# EMS-based mutants are useful for enhancing drought tolerance in spring wheat

**DOI:** 10.1101/2021.01.05.425390

**Authors:** Sadaf Zahra, Sana Zulfiqar, Momina Hussain, Muhammad Akhtar, Tayyaba Shaheen, Mehboob-ur-Rahman

## Abstract

Sustainable wheat production in drought prone areas can be achieved by developing resilient wheat varieties. In the present study, chemical mutagenesis was used to induce mutations in a cultivated wheat variety ‘NN-Gandum-1’. In total, 44 mutants were selected based on their high yield potential for exposing to well-watered (W_1_) and rainfed (W_2_) conditions for one season. Then 24 mutants were selected, and were exposed to W_1_ and W_2_ regimes. On the basis of least relative reduction in physiological parameters under W_2_ regime, five mutants were selected for conducting exome capturing assays. In total, 184 SNPs were identified in nine genes (ABC transporter type 1, Aspartic peptidase, Cytochrome P450, transmembrane domain, Heavy metal-associated domain, HMA, NAC domain, NAD (P)-binding domain, S-type anion channel, Ubiquitin-conjugating enzyme E2 and UDP-glucuronosyl/UDP-glucosyltransferase). Maximum number of mutations were observed in chr.2D, which contained mutations in three genes i.e. ABC transporter type 1, NAD (P)-binding domain and UDP-glucuronosyl/UDP-glucosyltransferase which may have a role in conferring drought tolerance. The selected mutants were further tested for studying their biochemical responses under both the regimes for two years. The extent of membrane damage was estimated through malondialdehydeand hydrogen per oxidase and tolerance to drought stress was assessed via antioxidant enzymes in leaves. The selected mutants under drought stress increased the accumulation of proline content, total soluble sugars, total free amino acids, while decreased total chlorophyll content, carotenoids and total soluble protein. Finally, the procedure of narrowing down the number of developed mutants from a large mutation population (>4000) is found useful for exploring the complex trait like drought without compromising yield potential. These mutants can further be explored to understand the genetic circuits of drought tolerance in wheat which will pave the way towards improving livelihood of resource poor farming community mostly relying on cereal food.

## Introduction

Wheat (*Triticum aestivum* L.) is cultivated in more than 60 countries on 219.52 million hectares with an annual production of 758.27 million metric tons (https://apps.fas.usda.gov/psdonline/circulars/production.pdf). Cereals have central role in global food security, predominantly in developed countries. Huge efforts have been made to improve/sustain wheat production by developing advanced varieties with improved genetics. However, there are still many factors which significantly decrease the wheat production [1].

Water deficit is predominant among the abiotic stresses that risks the cultivation of cereal crops [2–3]. Around 60% of wheat production is primarily affected by the limited supply of water [4–6]. Under the current scenario of climatic change, it could be inferred that water stress will aggravate in future [7–8]. Currently, improving yield potential and breeding drought tolerant wheat varieties are in progress to meet the increasing food demand of growing population of the world [9–12].With the onset of first green revolution (thirst agriculture revolution) between 1950s and 1960s, substantial advancements towards increasing wheat production were made by introducing dwarfing genes into the old wheat cultivars using conventional breeding techniques [13]. This improved germplasm containing the dwarfing genes was widely used for developing new wheat varieties, as a result, the wheat production increased significantly. There are still many areas left undiscovered which together reduce the wheat production globally especially drought stress [14–15]. To survive water stress via conventional breeding is challenging as it is difficult to regulate the multigenic response of abiotic stress tolerance [16–17].To combat dehydration, plants bring several changes in their physiological, morphological, biochemical and molecular mechanisms [18–21].

Understanding the adaptive response of important phenotypic traits that contribute to improved productivity during stress is essential in order to comprehend physiological as well as genetic methods of wheat adaptation [22–23]. As a response mechanism, some biochemical catalysts are produced, these alter the expression and regulation of multiple osmolytes, consequently leading to adverse homeostasis disruption [24] by affecting plant cellular mechanisms [25–26]. Drought-induced production of chemically reactive oxygen species (ROS) that expose cells to oxidative damage, results in membrane disability (lipid peroxidation), leakage of ions, breakage or cleavage of DNA strand at various levels, degradation of biomolecules, eventually leading to tissue damage and programmed cell death [27–30]. Though, cereals under abiotic stresses accumulate low molecular weight compatible solutes like sugars and proline to improve tolerance to drought stress [31]. While, total chlorophyll and carotenoid contents decline under water restricted conditions [32].

Various anti-oxidants (both enzymatic and non-enzymatic) play important role in plants to combat dehydration by controlling oxidative damage [33]. It was observed that concentration of various antioxidant enzymes like peroxidase (POD), superoxide dismutase (SOD) and catalase (CAT) increased significantly to mediate the effects of ROS for mitigating the oxidative stress in wheat [34–37]. Hence, enzymatic antioxidants have important role in coping drought stress, and thus can be used in screening drought tolerant wheat varieties [10].

For improving the genetics of wheat cultivars for coping drought, it is important to create allelic diversity followed by screening the wheat material under rainfed conditions. Mutagenesis had been an effective strategy to develop improved wheat genotypes and cope drought stress [38–41]. It was widely used to induce genetic variations and also the most effective and economical means for combating stresses [42]. Mutations can be spontaneous or induced artificially by exposing the biological material with physical or chemical mutagens [43]. Either for forward or reverse genetics, chemical mutagenesis is a standard choice for analysing gene function [44]. EMS (ethyl methane sulphonate), is extensively applied chemical mutagen which creates abrupt point mutations in plant genome [45–47]. Resultantly, mutated wheat lines and dwarf mutants have been produced, which are characterized to be drought resistant [48–49].

With the genetic advancements, NGS (next-generation sequencing) tools can be used to assess genetic variation and polymorphism [50–51]. Like other polyploids, wheat genome can withstand neutral mutations as these accumulated changes in genes which have no significant effect on survival of wheat plant [52–53]. For many plant species, complement of variant can be mapped via WGS (whole genome sequencing) [54] and these causal mutations can be recognized by identifying mapping regions [55]. Wheat has large genome size of around 17.6 GB and sequencing the whole wheat genome is quite expensive (International Wheat Genome Sequencing Consortium, 2014). In fact, exome-capturing approach can be used to lessen the cost of WGS [56]. Exome-capturing has been used to detect and sequence genetic variants among coding sequence (CDS) within wheat genome [57] and SNPs discovered by the exome capture assay can be helpful to detect mutations in putative candidate genes, mapping markers could also be developed [58, 59].

In the present investigation, five mutants (NN1-M-363, NN1-M-506, NN1-M-700, NN1-M-701 and NN1-M-1621) were selected from large mutant population of a wheat cultivar NN-Gandum-1 (NN-1). These mutants were subjected to exome capturing to detect SNPs relevant to drought resistance during M_5_ generation. The genetic diversity among wheat genotypes and drought tolerant mutant lines were further studied by applying different biochemical parameters for two years (M_6_ and M_7_ generation). The information generated through these studies can be used for exploring the genetic circuits of drought tolerance in wheat which will help in improving drought tolerant wheat cultivars.

## Material and Methods

### Experimental Growth Conditions

A high yielding wheat variety ‘NN-Gandum-1’, developed by the Plant Genomics and Molecular Breeding (PGMB) Lab, National Institute for Biotechnology and Genetic Engineering (NIBGE), Faisalabad, Pak, was selected for developing mutant population. The seed of this cultivar was exposed to an optimized concentration of 0.8% (v/v) of ethyl methane sulphonate for 2 h at 35°C. The seed were sterilized by performing three washings with 5% sodium hypochlorite and 70% ethanol to eradicate their residues. The procedure for developing the mutant population was described earlier [47].

### Generation advancement and screening /selection of mutant lines

The mutated population was advanced to M_4_ generation by raising single spike rows of M_1_ through M_3_. From planting to harvesting, recommended agronomic practices were applied to each generation. In total, 44 mutant lines were selected on the basis of their yield response and grown in randomized complete block design (RCBD) in three replicates under two water regimes i.e. well-watered conditions (W_1_ regimes) i.e. four irrigations while other set was grown in rainfed conditions (W_2_ regimes) i.e. ~170 mm rainfall. In total, 24 high yielding mutant lines were selected on the basis of drought tolerance indices and advanced to M_5_ generation. The drought indices such as tolerance index (TOL), geometric mean productivity (GMP), mean productivity (MP), stress susceptibility index (SSI) and stress tolerance index (STI) were calculated by following formulas:

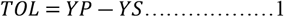

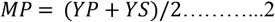

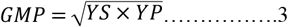

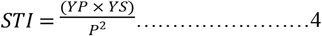

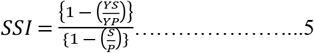

Here, Ys and Yp represent grain yield under drought stress and normal condition respectively. Mean yield of all genotypes under normal condition is denoted as P and S indicates the mean yield of genotypes under stress.

### Physiological parameters

These 24 mutant lines along with NN-1 (wild) were tested for physiological parameters such as photosynthetic absorbance rate (PAR) at leaf surface, sub-stomatal conductance, transpiration and photosynthetic rate under both the water regimes (W_1_ and W_2_ regime). Data was recorded using LcPro-SD portable Photosynthetic System. A total of five mutant lines (NN1-M-363, NN1-M-506, NN1-M-700, NN1-M-701 and NN1-M-1621) were selected on the basis of physiological parameters (Fig. 1).

**Fig 1.**
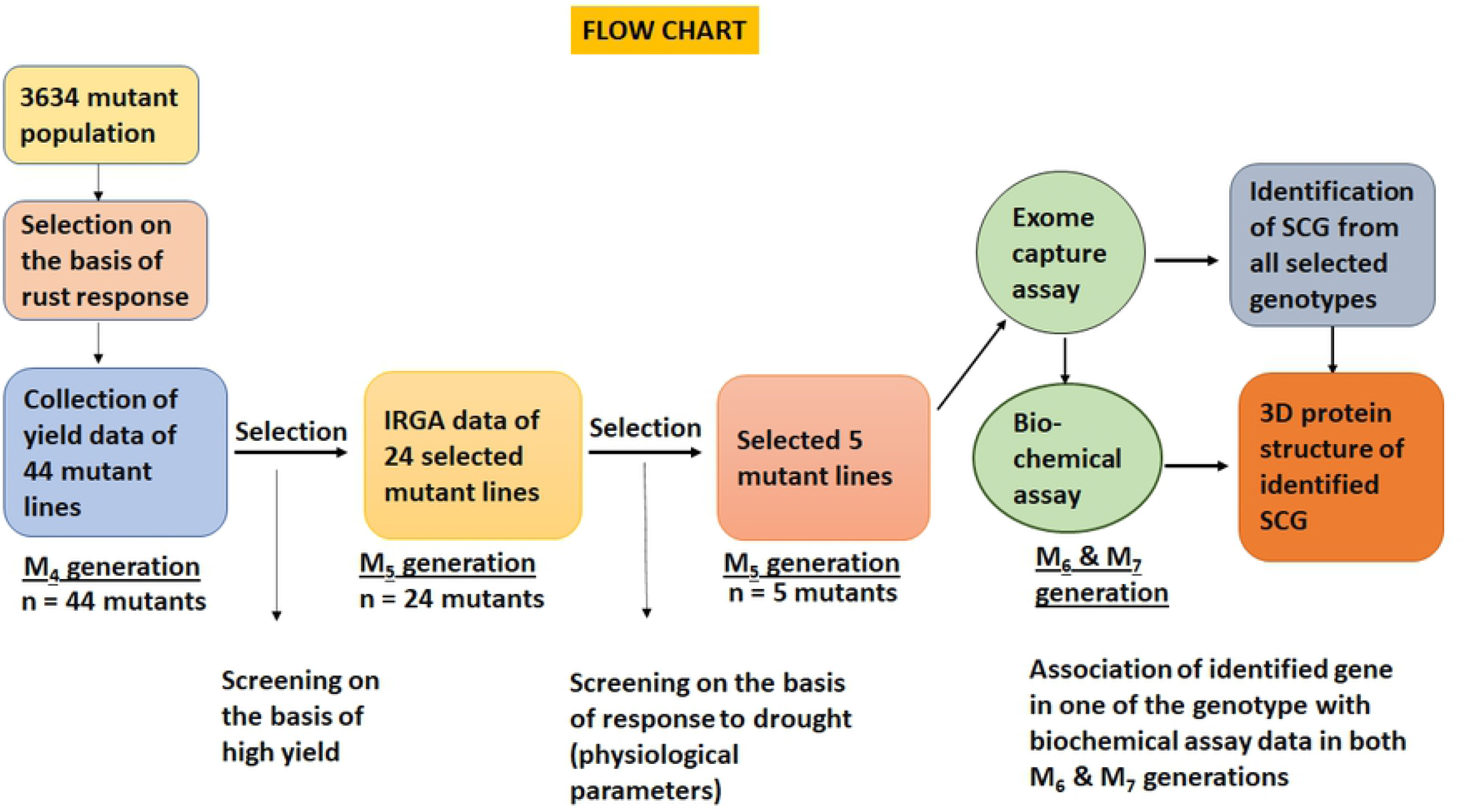
Flow chart showing the narrow down strategy for the identification.

### Exome Capture assay

These selected mutants and their parent NN-1 (wild) were sown for isolation of a high quality DNA by adopting the protocol described by [60] with few amendments. Genomic DNA concentration was measured with Nanodrop and normalized to 200ng /μl by adding (0.1 mM) EDTA, (10mM) Tris-HCl, maintained at pH 8 and making solution of 130 μl for every sample.

### Preparation of wheat exome captures libraries

The genomic DNA from individual samples were fragmented using Covaris E220 machine (UCD genome centre) for achieving the average fragment size (250-450 bp). The fragmented DNA was end repaired using End Repair enzyme, after that A-tail and adapter was added to these DNA fragments using a KAPA’s kit and Bioo Adapters (Sciclone G3 robot). The libraries with adapter ligation were exposed to Ampure beads for the size selection as per the selection protocol given in KAPA kits. Before hybridization, amplification step was completed for pre-capture of libraries with standard Nimblegen procedure. For each capture, five PCR cycles were performed. In total, 150 ng of each library product and oligoes for each capture were pooled to make 1.2 μg of total DNA for conducting hybridization. In a thermal cycler, hybridization was done for 68-72 hours outside of the robot (hybridization temperature was constant at 47°C and the lid was programmed for 57°C). Then washed and recovered the captured hybrids. Using post capture primers, PCR was done for amplification of these captures. However, for quality check qPCR, Bioanalyzer was operated. Finally, for sequencing the pooled and final captured libraries with Illumina Hi-Seq 2000 (Illumina) were submitted to Beijing Genome Institute (BGI).

### Analysis and arrangement of mutant’s reads against the wheat draft genome

The sequence of each DNA fragment from start to stop codon was assessed using bioinformatic tools like Fastq software, ‘bwaaln’ and ‘bwa sampe’ programs, samtools and bamtools, MAPS (http://comailab.genomecenter.ucdavis.edu/index.php/MAPS) and ‘mpileup’ pipeline (http://comailab.genomecenter.ucdavis.edu/index.php/Mpileup). The real SNPs were differentiated from mutations by applying an additional MAPS feature. The homozygous and heterozygous threshold was setup independently. Using Ensembl Variant Effect Predictor (VEP) release 78 in offline mode, the effect of mutation on gene function were predicted and detected.

### Biochemical Assays

For assessment of drought resistance, selected mutants were further tested for studying their biochemical responses under both the regimes (W_1_ and W_2_) for two consecutive normal wheat growing seasons.

Various biochemical parameters such as total soluble proteins (TSPs), total soluble sugars (TSSs), total free amino acids (TFAs), total chlorophyll contents, carotenoid contents, proline contents, enzymatic and non-enzymatic assays were analysed on M_6_ and M_7_ generations. For carrying out the aforementioned assays, fresh leaf samples were collected from the selected mutants and wild type in ice. The total TSPs (mg g^-1^ FW) were measured by following a published method [61]. In total, 0.5 g leaf tissues were homogenized into 10 ml phosphate buffer (0.2 M) and centrifuged at 4000 rpm for five min. After centrifugation, 1 ml supernatant was added into alkaline solution (1mL) and then 0.5 ml Folin-Phenol reagent (1:1 diluted) was added into the solution followed by the incubation of 30 min at room temperature. The optical density of the solution was measured at 620 nm using a spectrophotometer. Finally, the concentration of TSPs was estimated by comparing with the standard curve produced at various concentrations of bovine serum albumin (BSA).

For the determination of TSSs (mg g^-1^ FW), a protocol demonstrated by [62] was adopted. In total, 100 mg leaf tissue was homogenized by adding 5 mL ethanol solution (80%) and then incubated for six hours at 60°C. A total volume of 6 mL anthrone reagent was added and the solution was boiled for 10 min. The absorbance of the solution was recorded at 625 nm using a spectrophotometer. Finally, the sugar concentration was determined by comparing with a standard curve generated at different concentrations of glucose [63].

The TFAs (mg g^-1^ FW) were determined by adopting an assay described earlier [64]. A total of 0.5 g leaf tissue was added into 0.2 M phosphate buffer. The reaction mixture was prepared by adding1 mL of each, leaf extract, pyridine (10 %), ninhydrin (2%) and incubated for 30 min. The absorbance was recorded at 570 nm using spectrophotometer and the concentration of TFAs was calculated by comparing with standard curve of leucine.

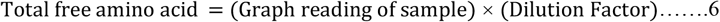

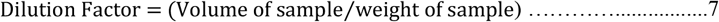

For the calculation of total chlorophyll and carotenoid contents, the protocols described by [65] and [66] were followed, respectively. In total, 0.5 g leaf tissue was homogenized into 5 ml acetone (80%) and the homogenate was centrifuged at 14000 rpm for five min. Eventually, the absorbance was monitored at 480, 645, 652 and 663 nm with the help of spectrophotometer. Through deploying the following formulas, chlorophyll (Chla, Chl_b_, Chl_t_) and carotenoids contents were calculated:

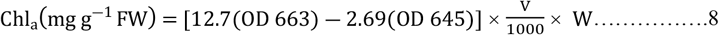

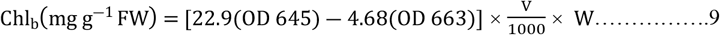

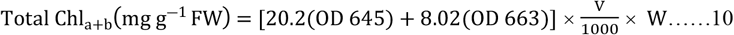

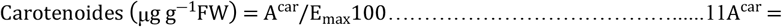

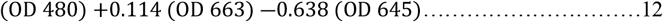

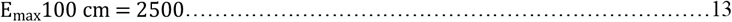

‘V’ is the volume of extract and ‘W’ is the weight of leaf sample

For the determination of proline contents (mmole g^-1^ FW), a published protocol was adopted [67]. A total of 0.5 g fresh leaf tissue was added into10 mL aqueous sulfo-salicylic acid (3%) and filtered. In total, 2 mL of the filtrate was added into the solution containing 2 mL each of acid ninhydrin solution and glacial acetic acid. The solution was then incubated at 100°C for one hour, and the absorbance was determined at 520 nm through spectrophotometer.

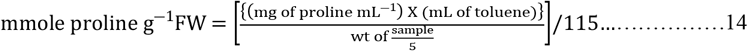

### Enzymatic Assays

Activity of superoxide dismutase (SOD, μ min^-1^mg^-1^) was measured by deploying a protocol demonstrated earlier [68]. The photochemical reduction of nitro blue tetrazolium (NBT) is inhibited by SOD at 560 nm and this inhibition is used to assay SOD activity. The reaction mixture was prepared by mixing 0.95 cm^3^ phosphate buffer (50 mM),0.5 cm^3^ methionine (13 mM), 1 cm^3^ NBT (50 μM), 0.5 cm^3^ EDTA (75 mM), 0.005 riboflavin, enzyme extract (50 μg mL)and ionized H_2_O (0.25 ml). The solution was irradiated by a fluorescent lamp (30V) for 15 min. Afterwards, inhibition in photochemical activity of NBT and non-irradiated reaction were analyzed at 560 nm, and were used to monitor the SOD activity.

The protocol devised by [68] was used to determine peroxidase activity (POD, min^-1^ g^-1^ FW). The reaction mixture consisted of phosphate buffer (50 mM), guaiacol (20 mM), H_2_O_2_ (40 mM) and enzyme extract (0.1 mL). POD activity was calculated by the estimation of the oxidation of guaiacol and peroxidation of H_2_O_2_ using an extinction coefficient of 2.47 mM−1 cm−1.

The catalase activity (CAT,min^-1^ g^-1^ FW) was measured by adopting a procedure described earlier [69]. The reaction mixture contained phosphate buffer (50 mM), H_2_O_2_ (5.9 mM) and 0.1 mL enzyme extract in a volume of 3 mL. The catalase activity was determined by the rate of H_2_O_2_ decomposition at 240 nm after every 20 sec.

The ascorbate peroxidase (APX, g^-1^ FW h^-1^) was quantified using the methodology proposed by [70]. The reaction mixture comprised of phosphate buffer (50 mM), sodium-EDTA (0.1 mM), H_2_O_2_ (12 mM), ascorbic acid (0.25 mM) and sample extract in a total volume of 1 ml. The rate of ascorbate oxidation at 470 nm was monitored after every 20 sec to measure APX activity. Finally, an extinction coefficient of 2.8 mM−1 cm−1 was used to calculate the APX activity.

### Non-Enzymatic Assays

For the detection of malondialdehyde contents (MDA, μmol g^-1^ FW), a published protocol was followed [71]. In total,1 g leaf tissue was added into 3cm^3^ of trichloroacetic acid solution (0.1%w/v) and centrifuged at 20000 rpm for 15 min. The cocktail was made by adding 0.5 cm^3^ supernatant and 3 cm^3^ thiobarbituric acid (0.5%) prepared in TCA (20%). Then mixture was incubated for one hour at 95°C and centrifuged at 10000 rpm for 10 min. Final concentration of MDA was estimated by measuring the change in optical density of supernatant at 532 nm and 600 nm using extinction coefficient (156 mmol^-1^cm^-1^).

The hydrogen per oxide concentration (H_2_O_2_, μmol g^-1^ FW) was measured by adopting a published protocol [67, 72]. In total, 0.2 g leaf tissue was mixed into trichloroacetic acid solution and centrifuged at 10000 rpm for 15 min. After centrifugation, a volume of 500 ul of each supernatant and potassium phosphate buffer (10 mM) were mixed together and added 1 mL of potassium iodide solution (1 M). The absorbance was recorded at 390 nm and activity of the H_2_O_2_ was determined by comparing with the standard curve drawn from H_2_O_2_ (Sigma-Aldrich) via spectrophotometer.

### Statistical analysis

Analysis of variance (ANOVA) and least significant difference (LSD) for each trait was estimated using Statistix 8.1 and SPSS16 software [73]. The statistical significance was computed at 5% probability.

## Results and Discussion

Wheat is dominantly a drought loving plant, and mitigates the water limited conditions by bringing changes in its morphological, physiological and biochemical properties. In the present experiment, selected wheat mutants were exposed to limited water conditions for studying their physiological as well as biochemical changes and their impact on yield.

### Productivity traits data

Increased grain yield of wheat is primary goal in drought affected areas. The use of drought resistant genotypes with higher yield is an effective approach to reduce harmful effects of drought [74]. In the present studies, significant differences were found for yield per unit area among the mutant lines grown under both the water regimes (well-watered, W_1_ and rain fed conditions, W_2_) at p<0.001. Wheat yield ranged from 3016.07 kg ha^-1^ to 5500.56 kg ha^-1^ in W_1_ regime. While in W_2_ regime, wheat yield fluctuated between 2256.32 and 4965.32 kg ha^-1^. Highest yield was depicted by NN1-M-451 with 9.73% relative reduction (RR) under W_2_ regime. While the lowest yield was determined for NN1-M-320 with 25.19% RR under W_2_ regime (Table 1). Similar results of wheat yield reduction under drought stress were reported earlier [75]. To identify high yielding drought tolerant genotypes under both the normal and drought conditions, drought tolerance indices was used as screening criteria [76]. In previous studies, geometric mean productivity (GMP), mean productivity (MP) and stress tolerance index (STI) were found to be the most appropriate indexes for the identification of drought tolerant cultivars [77–79].

**Table 1.** Yield and yield-drought related indices of 44 mutant lines along with wild type NN-1 sown under W_1_ and W_2_ regime during M_4_ generation

| Genotypes | Yield |  |  | Drought Tolerance Indices |  |  |  |  |  |
| --- | --- | --- | --- | --- | --- | --- | --- | --- | --- |
| Unit | W<sub>1</sub><br>regime | W<sub>2</sub><br>regime | RR | Tolerance<br>Index | Mean<br>Productivity | Geometric<br>Mean<br>Productivity | Stress<br>Tolerance<br>Index | Stress<br>Index | Stress<br>Susceptibility<br>Index |
| Conditions | kg ha-1 | kg ha-1 | % | TOL =<br>YP – YS | MP = (YP +<br>YS) / 2 | GMP =<br>SQRT of<br>(YS × YP) | STI =<br>(YP ×<br>YS) / (P)<br>2 | SI=1<br>– (YS<br>/ YP) | SSI = 1 – (YS<br>/ YP) / 1 – (S<br>/ P) |
| Local Check-1 | 3639.47 | 2956.32 | -18.77 | 683.15 | 3297.895 | 3280.158 | 0.6121 | 0.1877 | 2.0553 |
| Local Check-2 | 4711.35 | 3767.29 | -20.04 | 944.06 | 4239.32 | 4212.959 | 1.0097 | 0.2004 | 2.1941 |
| NN1-M-1 | 3760.56 | 3554.26 | -5.49 | 206.3 | 3657.41 | 3655.955 | 0.7604 | 0.0549 | 0.6007 |
| NN1-M-116 | 4359.29 | 4161.95 | -4.53 | 197.34 | 4260.62 | 4259.477 | 1.0321 | 0.0453 | 0.4957 |
| NN1-M-118 | 4179.89 | 3708.98 | -11.27 | 470.91 | 3944.435 | 3937.401 | 0.8819 | 0.1127 | 1.2336 |
| NN1-M-153 | 5224.87 | 4865.32 | -6.88 | 359.55 | 5045.095 | 5041.891 | 1.4461 | 0.0688 | 0.7535 |
| NN1-M-158 | 4280.8 | 3537.44 | -17.37 | 743.36 | 3909.12 | 3891.410 | 0.8615 | 0.1736 | 1.9014 |
| NN1-M-1621 | 3682.07 | 3558.74 | -3.35 | 123.33 | 3620.405 | 3619.880 | 0.7454 | 0.0335 | 0.3668 |
| NN1-M-205 | 3285.16 | 3130.44 | -4.71 | 154.72 | 3207.8 | 3206.867 | 0.5850 | 0.0471 | 0.5157 |
| NN1-M-2252 | 4585.77 | 4152.35 | -9.45 | 433.42 | 4369.06 | 4363.682 | 1.0832 | 0.0945 | 1.0349 |
| NN1-M-236 | 4366.02 | 4043.11 | -7.4 | 322.91 | 4204.565 | 4201.464 | 1.0042 | 0.0740 | 0.8098 |
| NN1-M-252 | 4004.98 | 3872.68 | -3.3 | 132.3 | 3938.83 | 3938.274 | 0.8823 | 0.0330 | 0.3617 |
| NN1-M-277 | 4466.93 | 3872.68 | -13.3 | 594.25 | 4169.805 | 4159.206 | 0.9841 | 0.1330 | 1.4566 |
| NN1-M-284 | 4058.8 | 3861.47 | -4.86 | 197.33 | 3960.135 | 3958.906 | 0.8916 | 0.0486 | 0.5323 |
| NN1-M-285 | 3426.44 | 2982.44 | -12.96 | 444 | 3204.44 | 3196.741 | 0.5813 | 0.1296 | 1.4188 |
| NN1-M-494 | 4314.44 | 3560.98 | -17.46 | 753.46 | 3937.71 | 3919.647 | 0.8740 | 0.1746 | 1.9122 |
| NN1-M-338 | 4747.23 | 4096.92 | -13.7 | 650.31 | 4422.075 | 4410.104 | 1.1064 | 0.1370 | 1.4999 |
| NN1-M-363 | 4415.35 | 4368.26 | -1.07 | 47.09 | 4391.805 | 4391.742 | 1.0972 | 0.0107 | 0.1168 |
| NN1-M-370 | 3482.5 | 3125.32 | -10.26 | 357.18 | 3303.91 | 3299.080 | 0.6192 | 0.1026 | 1.1230 |
| NN1-M-41 | 3996.01 | 3899.59 | -2.41 | 96.42 | 3947.8 | 3947.506 | 0.8865 | 0.0241 | 0.2642 |
| NN1-M-411 | 4594.74 | 4415.35 | -3.9 | 179.39 | 4505.045 | 4504.152 | 1.1541 | 0.0390 | 0.4275 |
| NN1-M-414 | 4500.56 | 4231.47 | -5.98 | 269.09 | 4366.015 | 4363.941 | 1.0834 | 0.0598 | 0.6547 |
| NN1-M-430 | 3825.59 | 3679.83 | -3.81 | 145.76 | 3752.71 | 3752.002 | 0.8008 | 0.0381 | 0.4172 |
| NN1-M-44 | 4789.84 | 4125.32 | -13.87 | 664.52 | 4457.58 | 4445.180 | 1.1241 | 0.1387 | 1.5191 |
| NN1-M-451 | 5500.56 | 4965.32 | -9.73 | 535.24 | 5232.94 | 5226.092 | 1.5537 | 0.0973 | 1.0655 |
| NN1-M-474 | 4588.02 | 3789.71 | -17.4 | 798.31 | 4188.865 | 4169.804 | 0.9891 | 0.1740 | 1.9052 |
| NN1-M-490 | 4419.83 | 4110.38 | -7 | 309.45 | 4265.105 | 4262.298 | 1.0335 | 0.0700 | 0.7666 |
| NN1-M-493 | 4601.47 | 3865.32 | -16 | 736.15 | 4233.395 | 4217.363 | 1.0118 | 0.1600 | 1.7517 |
| NN1-M-320 | 3016.07 | 2256.32 | -25.19 | 759.75 | 2636.195 | 2608.681 | 0.3871 | 0.2519 | 2.7582 |
| NN1-M-504 | 4386.62 | 3856.32 | -7.89 | 530.3 | 4121.47 | 4112.932 | 0.9623 | 0.1209 | 1.3237 |
| NN1-M-506 | 4612.68 | 4487.11 | -2.72 | 125.57 | 4549.895 | 4549.462 | 1.1774 | 0.0272 | 0.2981 |
| NN1-M-514 | 3330.01 | 2937.59 | -11.78 | 392.42 | 3133.8 | 3127.652 | 0.5565 | 0.1178 | 1.2903 |
| NN1-M-535 | 4330.14 | 3774.01 | -12.84 | 556.13 | 4052.075 | 4042.523 | 0.9297 | 0.1284 | 1.4063 |
| NN1-M-537 | 4610.44 | 3861.47 | -16.25 | 748.97 | 4235.955 | 4219.369 | 1.0128 | 0.1625 | 1.7788 |
| NN1-M-60 | 3648.44 | 3392.8 | -7.01 | 255.64 | 3520.62 | 3518.299 | 0.7042 | 0.0701 | 0.7672 |
| NN1-M-61 | 3569.95 | 3415.22 | -4.33 | 154.73 | 3492.585 | 3491.728 | 0.6936 | 0.0433 | 0.4746 |
| NN1-M-628 | 4321.17 | 4231.47 | -2.08 | 89.7 | 4276.32 | 4276.085 | 1.0402 | 0.0208 | 0.2273 |
| NN1-M-700 | 4224.74 | 4105.89 | -2.81 | 118.85 | 4165.315 | 4164.891 | 0.9868 | 0.0281 | 0.3080 |
| NN1-M-701 | 4704.62 | 4566.86 | -2.93 | 137.76 | 4635.74 | 4635.228 | 1.2222 | 0.0293 | 0.3206 |
| NN1-M-764 | 3993.77 | 3823.35 | -4.27 | 170.42 | 3908.56 | 3907.631 | 0.8686 | 0.0427 | 0.4672 |
| NN1-M-827 | 4608.2 | 4361.53 | -5.35 | 246.67 | 4484.865 | 4483.169 | 1.1434 | 0.0535 | 0.5861 |
| NN1-M-83 | 4027.41 | 3836.74 | -4.73 | 190.67 | 3932.075 | 3930.919 | 0.8790 | 0.0473 | 0.5184 |
| NN1-M-85 | 4099.17 | 3832.32 | -6.51 | 266.85 | 3965.745 | 3963.500 | 0.8937 | 0.0651 | 0.7128 |
| NN1-M-852 | 4253.89 | 3836.8 | -9.8 | 417.09 | 4045.345 | 4039.966 | 0.9285 | 0.0980 | 1.0736 |
| NN1-M-88 | 4191.11 | 3256.32 | -22.3 | 934.79 | 3723.715 | 3694.265 | 0.7764 | 0.2230 | 2.4422 |
| NN1-M-882 | 3269.47 | 2865.32 | -12.36 | 404.15 | 3067.395 | 3060.732 | 0.5329 | 0.1236 | 1.3535 |
| NN1-M-parent | 4249.41 | 4102.32 | -3.46 | 147.09 | 4175.865 | 4175.217 | 0.9917 | 0.0346 | 0.3790 |
| Mean | 4192.68 | 3809.77 | -9.13 | 387.167 | 4003.349 | 3996.499 | 0.9230 | 0.0926 | 1.0136 |
| Max | 5500.56 | 4965.32 | -9.73 | 944.060 | 5232.940 | 5226.092 | 1.5537 | 0.2519 | 2.7582 |
| Min | 3016.07 | 2256.32 | -25.19 | 47.090 | 2636.195 | 2608.681 | 0.3871 | 0.0107 | 0.1168 |
| SD | 514.26 | 530.17 | 3.09 | 253.390 | 507.070 | 508.701 | 0.2273 | 0.0606 | 0.6635 |
| SE | 75.074 | 77.398 | 3.09 | 36.961 | 73.964 | 74.202 | 0.0332 | 0.0088 | 0.0968 |

It was reported earlier that genotypes having minimum tolerance index (TOL) and stress susceptibility index (SSI) values were least sensitive to drought and selection exclusively based upon these indices directs high yielding genotypes under drought conditions [5–75, 80]. In this study minimum TOL was calculated for NN1-M-363, NN1-M-628, NN1-M-41, NN1-M-700, NN1-M-1621 and NN1-M-506. While for SSI, NN1-M-363, NN1-M-628, NN1-M-41, NN1-M-506, NN1-M-700 and NN1-M-701 depicted lowest values (Table 1). Under normal conditions, these genotypes demonstrated high yield while, under W_2_ regime their yield was reduced. The importance of the SSI was explained by [81] which and demonstrated that genotypes with less than one SSI value were tolerant to drought.

We selected 24 mutant lines which showed high MP and GMP values and minimum values for SSI and TOL as these were drought resistance and high yielding genotypes.

### Physiological parameters analysis

Photosynthesis, a primary metabolic process which determines crop production is affected by drought stress [82]. In the current investigation, a total of 24 mutant lines having high yield were selected from M_4_ population. These mutant lines along with wild type were tested for physiological parameters such as photosynthetically active radiation (PAR) at leaf surface, sub-stomatal conductance (g_s_), transpiration rate (E) and net photosynthetic rate (Pn) under W_1_ and W_2_ regimes during M_5_ generation. Significant variation were found for all the physiological parameters among all the mutant lines under both water regimes at p<0.001. Water stress caused prominent decrease in net photosynthetic rate (Pn), sub-stomatal conductance (g_s_) and transpiration rate (E) in wheat and different crops [34, 83]. Under drought stress plant immediately closes stomata to maintain cellular moisture level [84–86]. Moreover, diffusion of CO_2_ from outside atmosphere to the sub-stomatal cavity is reduced, resultantly decrease the stomatal conductance (g_s_), and is the major reason of reduction in the photosynthetic rate (Pn) during drought [84]. Some other studies have also reported reduction in net photosynthetic rate and sub-stomatal conductance in important crops during drought condition, such as in rice [87] and wheat [88].

In the present study, least relative reduction in PAR at leaf surface was observed for NN1-M-700 (4.2%), NN1-M-701 (4.5%), NN1-M-1621 (4.6%) and NN1-M-363(5%) as compared with wild type (5.1%) under W_2_ regime (Fig 2). Regarding, sub-stomatal conductance minimum percentage reduction was recorded for NN1-M-506 (5.5%), NN1-M-1621 (6.5%), NN1-M-363 (7.2%), and NN1-M-701 (7.8%) compared to wild type (7.7%) during water deficient condition. Similar findings of drought induced reduction were observed in stomatal conductance [89]. In case of transpiration rate a decrease of 4.6% (NN1-M-363), 4.7% (NN1-M-506), 5.1% (wild type) and 5.5% (NN1-M-700) was demonstrated. Reduction in transpiration rate were also reported in some other studies [83]. As far net photosynthetic rate is concerned, the mutants NN1-M-363 (10.93%), NN1-M-701 (11.14%) and NN1-M-700 (11.72%) showed minimum percentage reduction (S1 Table). Reduced inhibition of net photosynthetic rate under limited water conditions is of great importance for drought tolerance [90].

**Fig 2.**
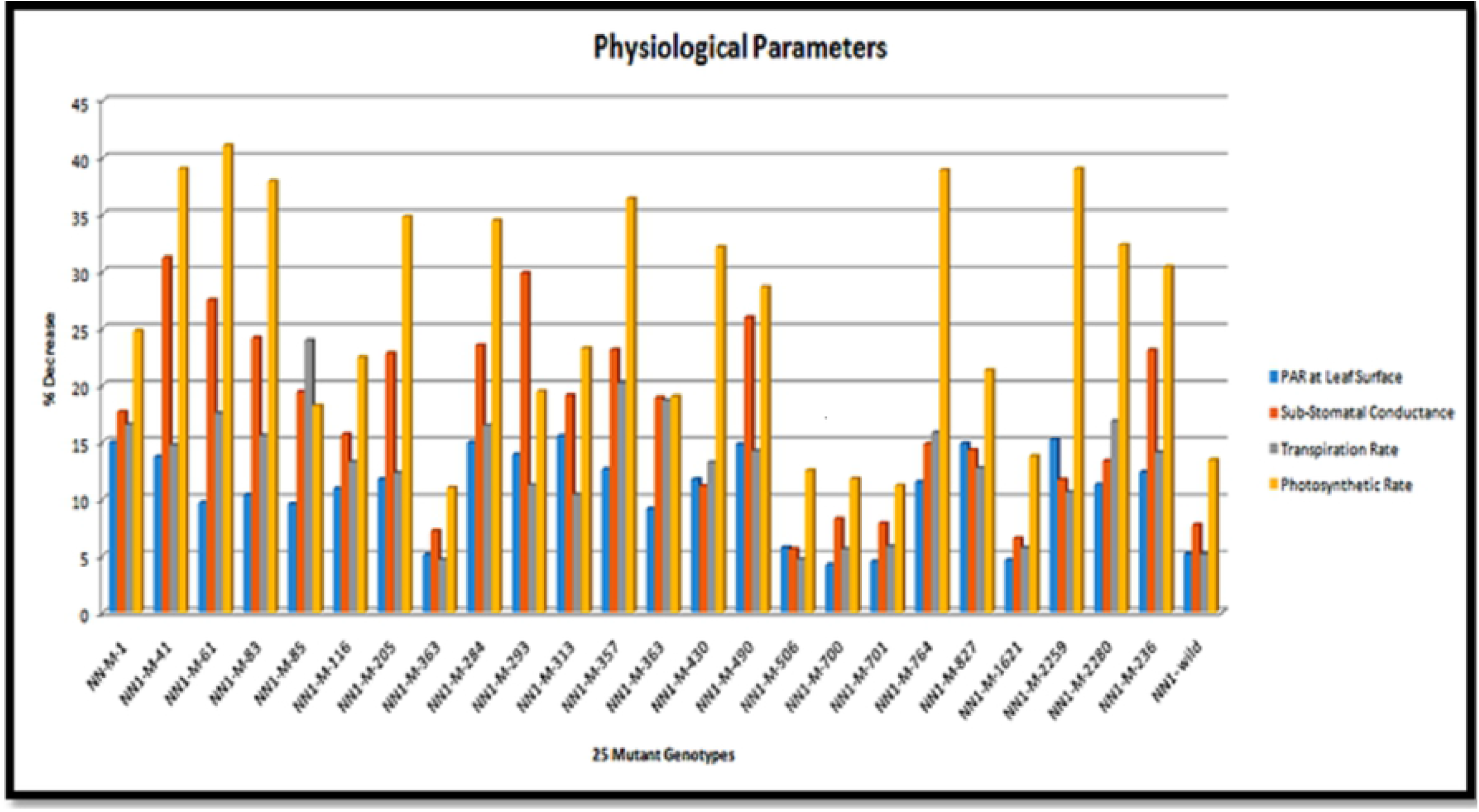
Identification of 5 mutants from 24 mutant lines on the base of physiological data. Physiological data: PAR at leaf surface, substomatal conductance, transpiration rate and photosynthetic rate

On the basis of physiological parameters, five mutant lines i.e. NN1-M-363, NN1-M-506, NN1-M-700, NN1-M-701 and NN1-M-1621 showing minimum reduction in PAR at leaf surface, net photosynthetic rate, sub-stomatal conductance and transpiration rate were selected for conducting exome capturing assays.

### Exome capture assay

The hexaploid wheat has relatively gigantic genome size of about 17.6 GB, and it is difficult to identify mutations in such a large genome through using whole genome sequencing but exome capturing is one of the most suitable assays which can be exploited to detect mutations induced by EMS. By deploying exome sequencing, mutations can be detected in coding regions of a gene (exon) only which can alter the gene function and expression. The exome sequencing can be exploited for searching mutations in expressed part of the genes using multiple software including MAPS, VEP, etc. It is well known that fluctuations in genomes size are contributed by the non-exonic portions of the genome, thus exome size remains about the same in all plant species. Consequently, exome capture can be practiced in various crops regardless of the genome size and cost involved in conducting exome capture assay. This assay is extremely handy and economical for studying the genome of those species where reference genome assembly has not been constructed yet. We selected five mutant lines i.e. NN1-M-363, NN1-M-506, NN1-M-700, NN1-M-701 and NN1-M-1621 on the basis of their response to drought. These mutants were exposed to exome capture assay for the identification of SNPs.

In total, 184 SNPs were identified in different genes which are involved in conferring drought tolerance. Overall, 96, 55, 14, 11 and 8 SNPs were detected in N1-M-701, NN1-M-700, NN1-M-506, NN1-M-1621 and NN1-M-363, respectively (Table 2). The maximum number of SNPs both heterozygous and homozygous were observed in chr-2D (31 SNPs), however, few SNPs were found in chr-5D (7 SNPs) and chr-7B (3 SNPs). These SNPs dispersed on various wheat chromosomes are depicted in a Circos plot (Fig 3).

**Fig 3.**
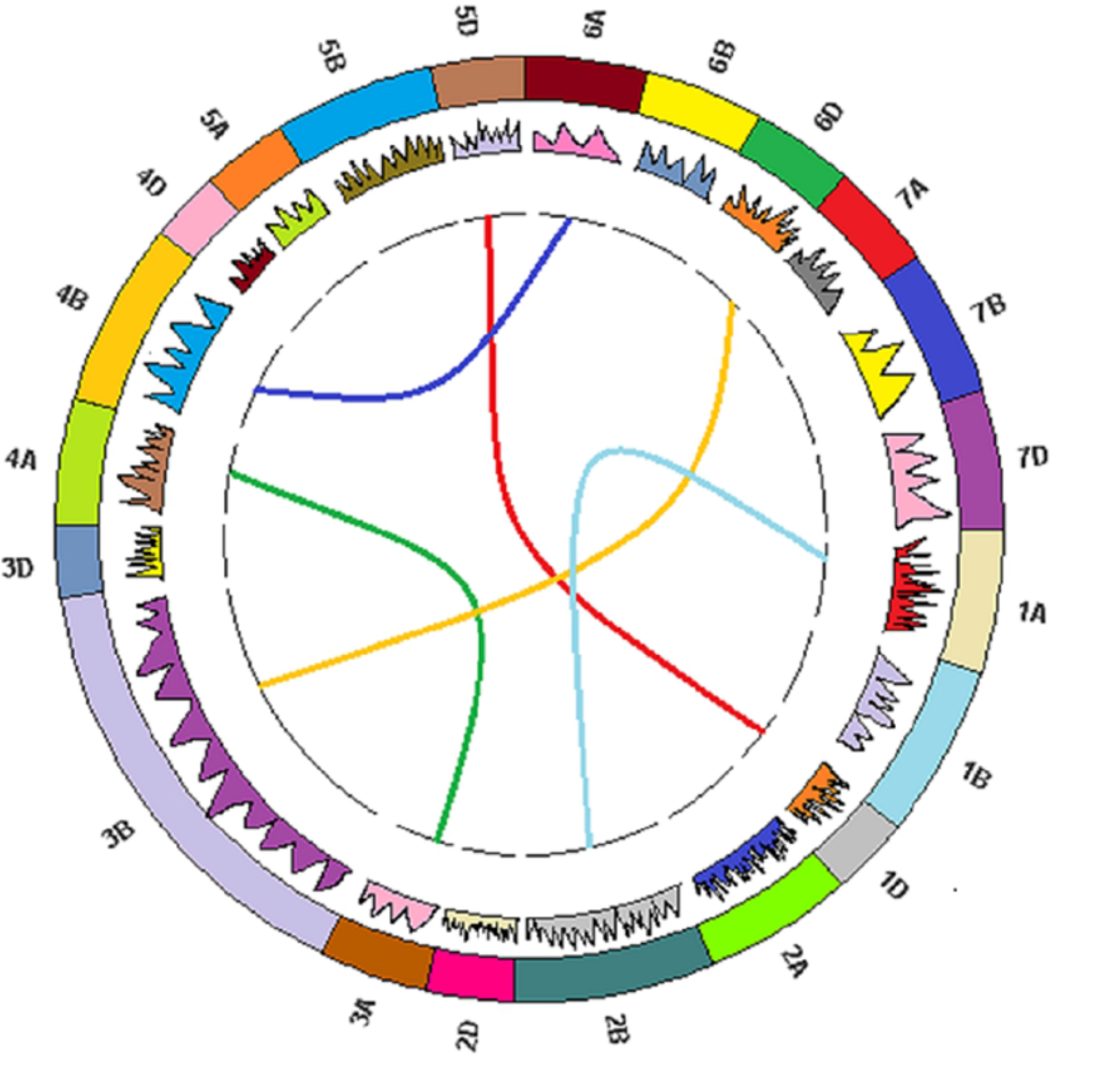
J-Circos Plot: First Circos plot indicates the size of exonic region of all wheat chromosome. Graph bar in second circle shows dispersion of 184 SNPs on different chromosomes of wheat genome (A, B & D). Lines in third circle indicates the distribution of SNPs on five wheat mutants i.e. Blue; NN1-M-363, Green; NN1-M-506, Yellow; NN1-M-700, Red; NN1-M-701 and Light blue; NN1-M-1621

**Table 2.**
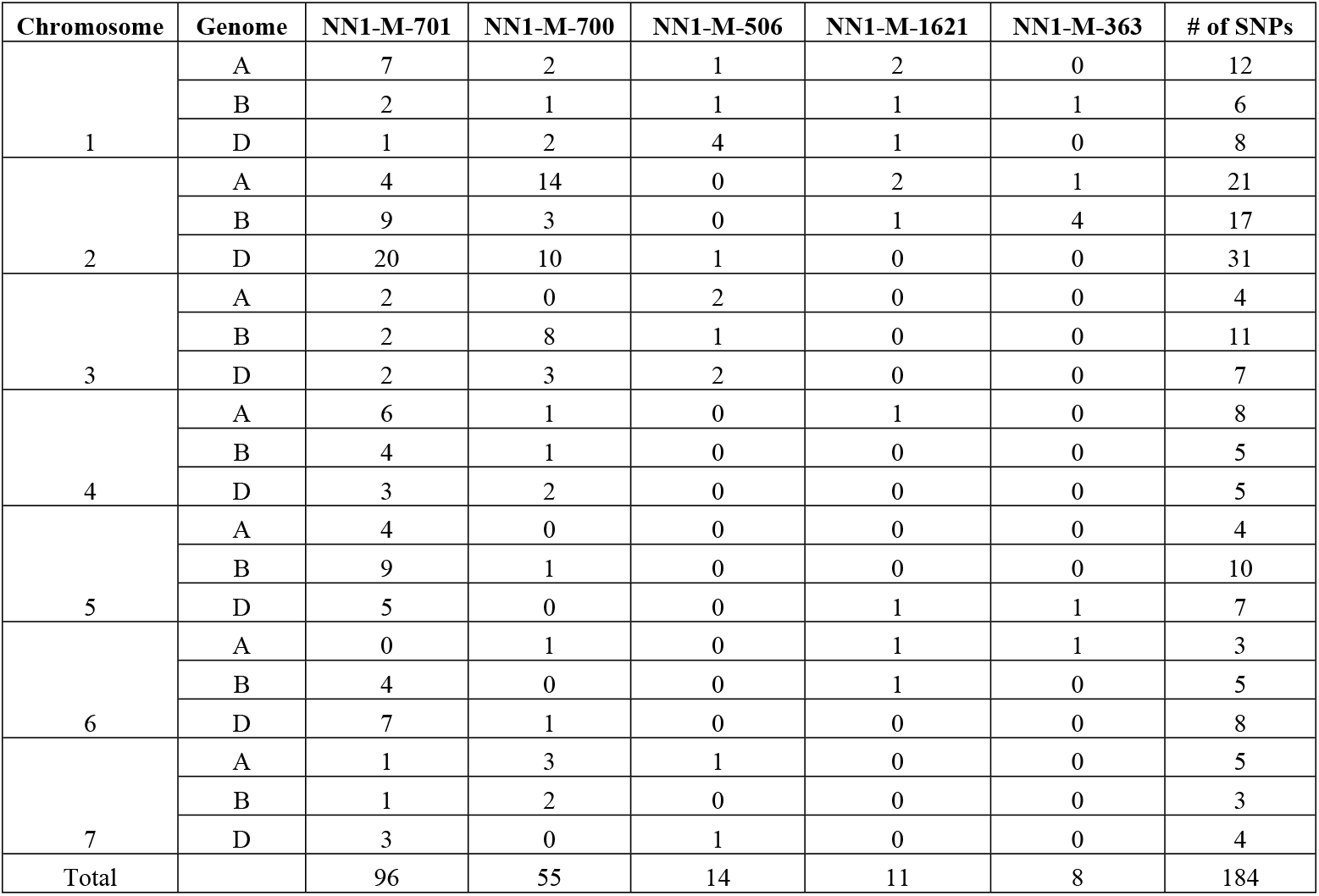
SNPs detected in the selected mutants

| Chromosome | Genome | NN1-M-701 | NN1-M-700 | NN1-M-506 | NN1-M-1621 | NN1-M-363 | # of SNPs |
| --- | --- | --- | --- | --- | --- | --- | --- |
| 1 | A | 7 | 2 | 1 | 2 | 0 | 12 |
|  | B | 2 | 1 | 1 | 1 | 1 | 6 |
|  | D | 1 | 2 | 4 | 1 | 0 | 8 |
| 2 | A | 4 | 14 | 0 | 2 | 1 | 21 |
|  | B | 9 | 3 | 0 | 1 | 4 | 17 |
|  | D | 20 | 10 | 1 | 0 | 0 | 31 |
| 3 | A | 2 | 0 | 2 | 0 | 0 | 4 |
|  | B | 2 | 8 | 1 | 0 | 0 | 11 |
|  | D | 2 | 3 | 2 | 0 | 0 | 7 |
| 4 | A | 6 | 1 | 0 | 1 | 0 | 8 |
|  | B | 4 | 1 | 0 | 0 | 0 | 5 |
|  | D | 3 | 2 | 0 | 0 | 0 | 5 |
| 5 | A | 4 | 0 | 0 | 0 | 0 | 4 |
|  | B | 9 | 1 | 0 | 0 | 0 | 10 |
|  | D | 5 | 0 | 0 | 1 | 1 | 7 |
| 6 | A | 0 | 1 | 0 | 1 | 1 | 3 |
|  | B | 4 | 0 | 0 | 1 | 0 | 5 |
|  | D | 7 | 1 | 0 | 0 | 0 | 8 |
| 7 | A | 1 | 3 | 1 | 0 | 0 | 5 |
|  | B | 1 | 2 | 0 | 0 | 0 | 3 |
|  | D | 3 | 0 | 1 | 0 | 0 | 4 |
| Total |  | 96 | 55 | 14 | 11 | 8 | 184 |

Though, chemical mutagenesis is a random process and many mutations are functionally silent. Also, the frequency of mutations in each site of the genome fluctuates substantially. Certainly, it was proved that if concentration of the mutagen remains same, the rate of mutation ranges from 1-20 mutations per Mb in diverse entities [91]. Yet, the total number of base pairs involved in the analysis varies by chromosome, followed by measuring the density of mutations. The maximum number of mutations were detected in chr.2D. However, minimum mutation density was identified in chr.6A, chr.5A, chr.7B and chr.7D. The mean mutation rate was 0.04 mutations/Mb (Table 3).

**Table 3.** Frequency of mutation / Mb

| **Chromosome** | **Sequenced exon size**<br>**in each chromosome**<br>**(Mb)** | **# of SNPs** | **Number**<br>**of SNPs /**<br>**Mb** |
| --- | --- | --- | --- |
| 1A | 248 | 12 | 0.05 |
| 1B | 295 | 6 | 0.02 |
| 1D | 135 | 8 | 0.06 |
| 2A | 255 | 21 | 0.08 |
| 2B | 345 | 17 | 0.05 |
| 2D | 150 | 31 | 0.21 |
| 3A | 185 | 4 | 0.02 |
| 3B | 124 | 11 | 0.09 |
| 3D | 216 | 7 | 0.03 |
| 4A | 317 | 8 | 0.03 |
| 4B | 121 | 5 | 0.04 |
| 4D | 148 | 5 | 0.03 |
| 5A | 274 | 4 | 0.01 |
| 5B | 162 | 10 | 0.06 |
| 5D | 208 | 7 | 0.03 |
| 6A | 204 | 3 | 0.01 |
| 6B | 177 | 5 | 0.03 |
| 6D | 182 | 8 | 0.04 |
| 7A | 252 | 5 | 0.02 |
| 7B | 238 | 3 | 0.01 |
| 7D | 774 | 4 | 0.01 |
| Mean mutation | 5010 | 184 | 0.04 |

The main reasons of these fluctuations in mutation density owing to the variations in penetrability of EMS within seeds and the differential ability of the cell to repair the damaged DNA as shown in previous studies [44]. In several other studies, variation in normal density of mutation such as 19.6 mutations per Mb, 20.1 mutations per Mb, 23 mutations per Mb and 33 mutations per Mb were noticed in wheat [52, 91, 92]. In total, 121 mutations were reported in waxy genes in wheat, including silent, missense together with knockout by studying 2,348 EMS-mutated M_2_ plants [93]. Similarly, in four important genes i.e. *LBP, COMIT1, HCT2*, and *4CL1*, SNPs were recognized. In TILLING population of polyploid wheat, the mutation frequency was one mutation per 17.6 kb to 34.4 kb mutations. However, in case of *T. monococcum* only one mutation per 90 kb was identified in waxy genes [94].

In the present studies, out of the 184 SNPs, 31 were induced in genes closely related to abiotic stresses particularly drought such as *ABC transporter type 1, aspartic peptidase, cytochrome P450, transmembrane domain, heavy metal-associated domain, HMA, NAC domain, NAD(P)-binding domain, s-type anion channel, ubiquitin-conjugating enzyme E2* and *UDP-glucuronosyl/UDP-glucosyltransferase* (Table 4).

**Table 4:** Gene Functions

| Genotypes | Genes | Functions | References |
| --- | --- | --- | --- |
| NN1-M-701 | ABC transporter type 1, transmembrane domain | Regulate physiological activities | ---- |
| NN1-M-1621 | Aspartic peptidase | Plant senescence, stress responses, programmed cell death and reproduction | Isaura and Carlos, 2004 |
| NN1-M-700 | Cytochrome P450 | Growth, development, protects plant from stresses, establishment of complex signaling webs | Li et al., 2010; Jun et al., 2015 |
| NN1-M-701 |  |  |  |
| NN1-M-700 | Heavy metal-associated domain, HMA | Heavy metal transport and detoxification, plays role during drought, cold and salt stress | Barth et al., 2009; Zschiesche et al., 2015 |
| NN1-M-700 | NAC domain | Regulate biological processes, biotic and abiotic stress responses | Zhao et al., 2015; Chen et al., 2018 |
| NN1-M-700 | NAD(P)-binding domain | Abiotic stresses, developmental processes | Hashida et al., 2009 |
| NN1-M-701 |  |  |  |
| NN1-M-1621 |  |  |  |
| NN1-M-363 |  |  |  |
| NN1-M-701 | S-type anion channel | Regulate drought stress responses | Murata et al., 2015 |
| NN1-M-701 | Ubiquitin-conjugating enzyme E2 | DNA repair, cell cycle regulation, plant growth and development | Hochstrasser, M. 2003; Gao et al., 2017 |
| NN1-M-700 | UDP-glucuronosyl/UDP-glucosyltransferase | Drought stress tolerance, seed germination | Liu et al., 2015 |
| NN1-M-701 |  |  |  |

One mutation was observed in each of the *ABC transporter type 1, transmembrane domain, Aspartic peptidase, heavy metal-associated domain, HMA, NAC domain, S-type anion channel* and *Ubiquitin-conjugating enzyme E2* gene located on chr-2D of NN1-M-701 (114.5 Mb position), chr-2B of NN1-M-1621 (225.91 Mb position), chr-1A of NN1-M-700 (3.28 Mb position), chr-7B of NN1-M-700 (250.85 Mb position), chr-1B of NN1-M-701 (99.74 Mb position) and chr-1A of NN1-M-701 (3.88 Mb position) (Table 5). Similarly, four mutations were observed in *Cytochrome P450* gene in NN1-M-700 and one in NN1-M-701. Likewise, four mutations in NN1-M-700, six in NN1-M-701, and one each in NN1-M-1621, and NN1-M-363 were observed in *NAD(P)-binding domain* gene. A total of six mutations in NN1-M-701 and two in NN1-M-700 were identified in UDP-glucuronosyl/UDP-glucosyltransferase gene (Table 5).

**Table 5.** SNPs detected in example gene

| Chromosome | Position (Mbs) | Genotypes | Protein position | Amino acids | Codons | Gene |
| --- | --- | --- | --- | --- | --- | --- |
| 2D | 114.50 | NN1-M-701 | 120 | A/T | Gcg/Acg | ABC transporter type 1, transmembrane domain |
| 2B | 225.91 | NN1-M-1621 | 116 | R/K | aGg/aAg | Aspartic peptidase |
| 2A | 27.95 | NN1-M-700 | 347 | W/* | tgG/tgA | Cytochrome P450 |
| 2A | 7.50 |  | 171 | R/K | aGg/aAg |  |
| 2A | 7.50 |  | 434 | S/N | aGt/aAt |  |
| 3B | 89.37 |  | 98 | G/S | Ggc/Agc |  |
| 5B | 114.32 | NN1-M-701 | 281 | R/H | cGc/cAc |  |
| 1A | 3.28 | NN1-M-700 | 88 | V/M | Gtg/Atg | Heavy metal-associated domain, HMA |
| 7B | 250.85 | NN1-M-700 | 283 | W/* | tgG/tgA | NAC domain |
| 1A | 2.18 | NN1-M-700 | 174 | G/S | Ggc/Agc | NAD(P)-binding domain |
| 2D | 9.90 |  | 76 | V/I | Gtt/Att |  |
| 2D | 41.48 |  | 17 | G/D | gGc/gAc |  |
| 3B | 9.98 |  | 252 | M/I | atG/atA |  |
| 1D | 2.22 | NN1-M-701 | 133 | A/T | Gca/Aca |  |
| 2D | 9.87 |  | 55 | A/T | Gca/Aca |  |
| 2D | 134.47 |  | 195 | C/Y | tGc/tAc |  |
| 4D | 65.06 |  | 740 | E/K | Gaa/Aaa |  |
| 4D | 65.06 |  | 780 | E/K | Gaa/Aaa |  |
| 6B | 2.44 |  | 115 | E/K | Gag/Aag |  |
| 2A | 14.18 | NN1-M-1621 | 949 | G/D | gGc/gAc |  |
| 6A | 4.39 | NN1-M-363 | 51 | A/T | Gca/Aca |  |
| 1B | 99.74 | NN1-M-701 | 161 | Q/* | caG/tgA | S-type anion channel |
| 1A | 3.88 | NN1-M-701 | 192 | G/D | gGt/gAt | Ubiquitin-conjugating enzyme E2 |
| 2A | 7.14 | NN1-M-700 | 258 | G/S | Ggc/Agc | UDP-glucuronosyl/UDP-glucosyltransferase |
| 2D | 3.84 |  | 2 | G/D | gGt/gAt |  |
| 2B | 9.40 | NN1-M-701 | 248 | G/R | Ggg/Agg |  |
| 2B | 274.25 |  | 166 | G/S | Ggc/Agc |  |
| 2D | 3.84 |  | 50 | R/H | cGc/cAc |  |
| 5B | 191.93 |  | 321 | E/K | Gag/Aag |  |
| 5D | 154.00 |  | 35 | D/N | Gac/Aac |  |
| 6D | 140.48 |  | 83 | G/D | gGc/gAc |  |

### 3D protein structure

The SNP identified in chimeric allele (heavy metal-associated domain, HMA) of a drought resistant mutant (NN1-M-700) was located in HMA domain of chr.1A at 3.28 Mb position. Through computational analysis, it was demonstrated that this SNP causes a substitution of valine with methionine, resulting in a predicted altered protein structure (Fig 4). This mutation, therefore, is a candidate for contributing to the resistance phenotype in the mutant line.

**Fig 4.**
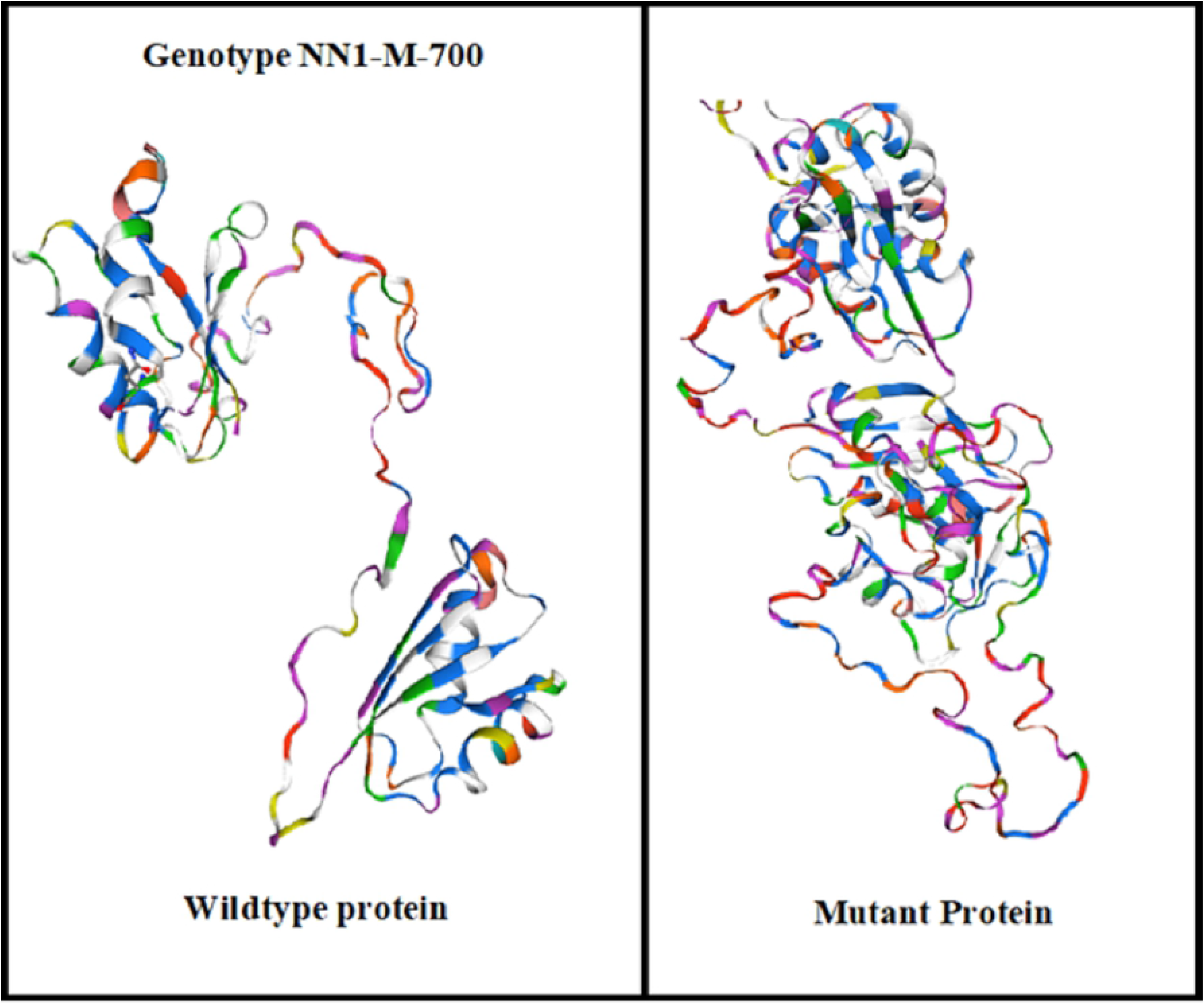
3D protein structure of single copy gene HMA identified in genotype NN1-M-700

### Biochemical assays

#### Total Soluble Proteins (TSPs)

The TSPs play a vital role in cellular dehydration tolerance in wheat and their accumulation in leaves helps in mitigating dehydration [95–96]. The current study indicated that drought stress significantly decreased the total soluble proteins in both generations (M_6_ and M_7_) of wheat. In M_6_ generation, NN1-M-363 and NN1-M-1621 exhibited 35 and 33% reduction in TSPs, respectively (S2 Table). While NN1-M-506 and NN1-M-701 showed less reduction in TSPs in both the generations under drought conditions (Fig 5). The decline in soluble proteins during dehydration might be linked with the decrease in protein biosynthesis [97]. In another study, soluble proteins were reduced after the exposure of drought stress in tomato and sorghum [98]. This indicates reduction in protein synthesis is due to its breakage to amino acids under water limited conditions. Contrary to the present study, synthesis of protein in plants was increased under stress condition regulated by endogenous cytokinins and eventually promoting drought tolerance [99].

**Fig 5.**
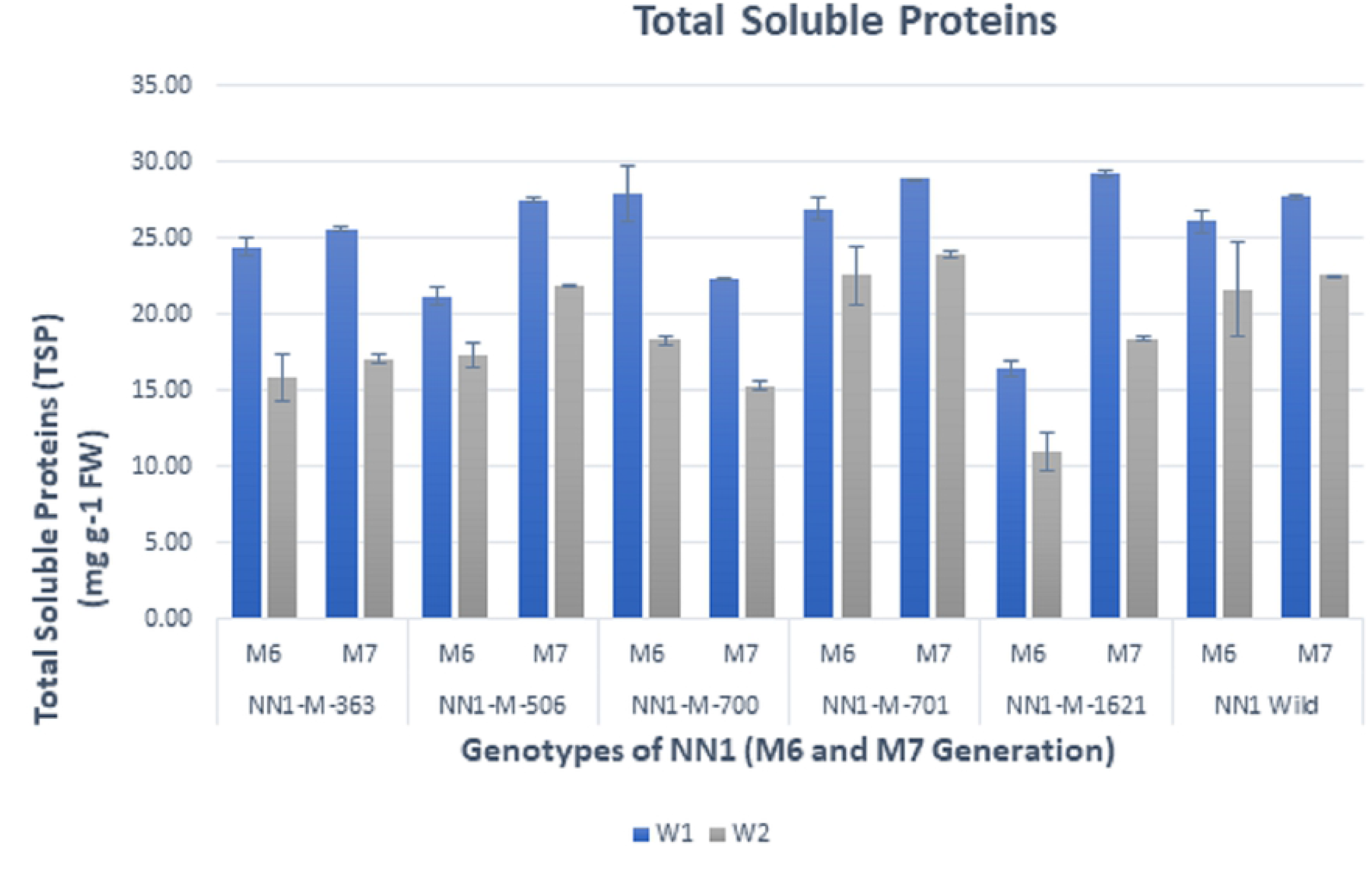
Total Soluble Proteins (TSPs) in NN-Gandum-1 genotypes. At two conditions: well-watered (control, W_1_ regime) and rainfed (stressed, W_2_ regime). Values presented are means ± S.D of three replicates per genotype in both M_6_ and M_7_ generation. Different color indicates stress condition, blue bar for W_1_-well watered (control) and grey bar for W_2_-rainfed (stressed) conditions

#### Total Soluble Sugars (TSSs)

Wheat plant accumulates high level of soluble sugars for mitigating water deficit. These sugars are synthesized during photosynthesis and act as substrate for carbon and energy metabolism in plants [100]. Soluble sugars work as osmoprotectants for leaf osmotic adjustment and protect the cells from dehydration [101]. A significant increase in soluble sugars in leaf during drought stress was observed in German chamomile (*Matricaria chamomilla* L.) [102]. In present investigation, soluble sugars have been increased by desiccation in both the generations (Fig 6). Wheat genotype NN1-M-701 showed highest increase, i.e. 64% and 69% in M_6_ and M_7_ generations, respectively (S2 Table). Such an increase in soluble sugar contents were also observed in common bermudagrass under limited water conditions [31].

**Fig 6.**
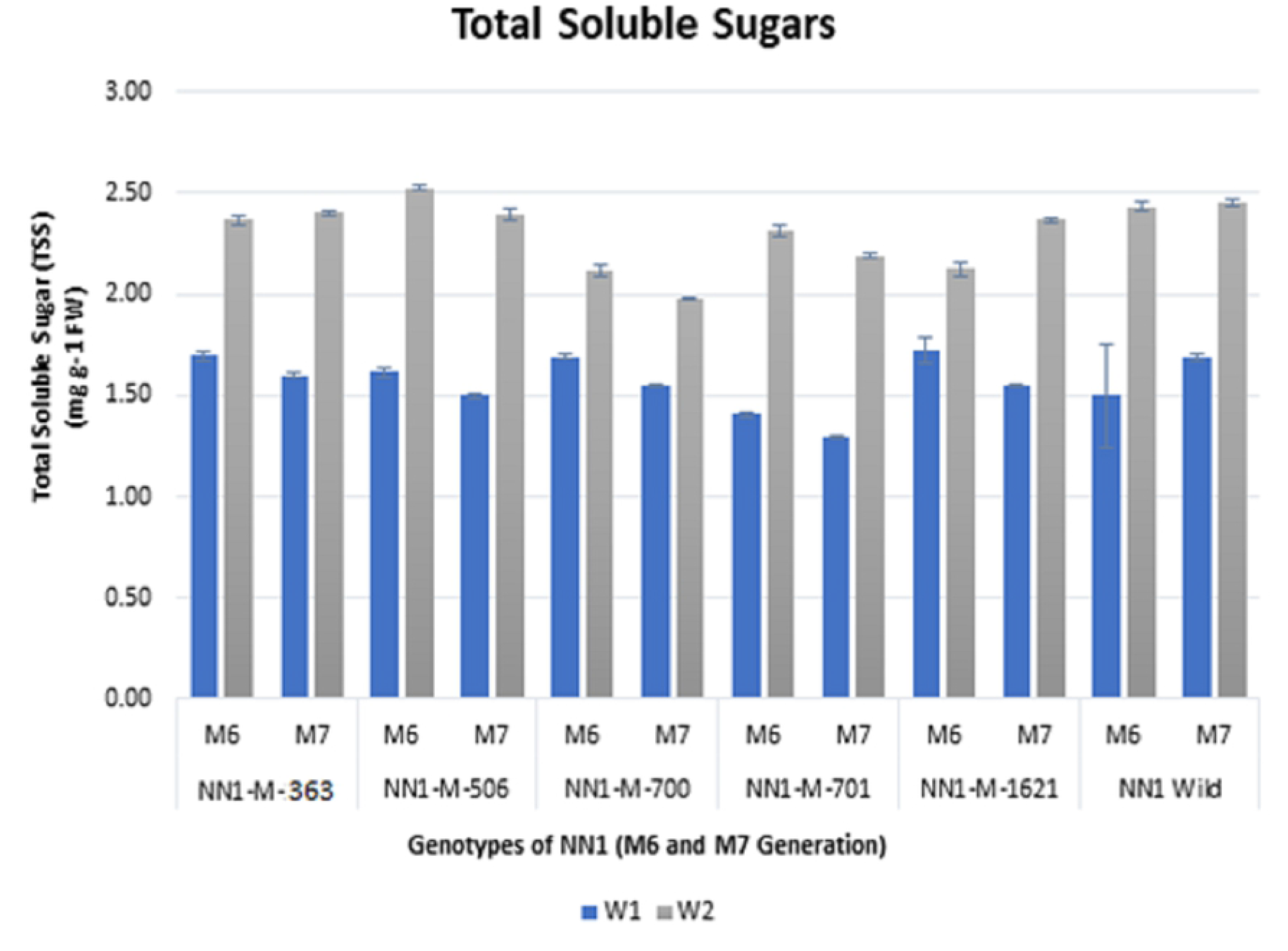
Total Soluble Sugars (TSSs) in NN-Gandum-1 genotype under well-watered (control, W_1_ regime) and rainfed (stressed, W_2_ regime) conditions. Values presented are means ± S.D of three replicates per genotype in both M_6_ and M_7_ generation. Different color indicates stress condition, blue bar for W_1_-well watered (control) and grey bar for W_2_-rainfed (stressed) conditions

#### Total Free Amino Acid (TFAs)

TFAs inhibit the production of reactive oxygen species (ROS) and regulate stomata, and defend macromolecules to maintain cellular water balance under drought stress. Free amino acids and sugars contents can be used as an essential selection criterion for drought tolerance [34]. In the current investigation, a remarkable increase in total free amino acid was recorded for all the mutants in both the generations (M_6_ and M_7_) (Fig 7). During M_6_ generation, 72% increase was demonstrated for NN1-M-506. While next year, 75% and 76% increase in TFAs were estimated for NN1-M-701 and NN1-M-506 respectively. Total free amino acids were accumulated more in M_7_ as compared to M_6_ generation. However, relatively less accumulation of free TFAs was observed in NN1-M-1621 under rainfed condition (Table S2). These results are in accordance with the earlier studies [103–104]. The increase in TFAs in present study indicates that stress tolerance in mutants resulted from the detoxification of ROS and perform functional role as compatible osmolyte.

**Fig 7.**
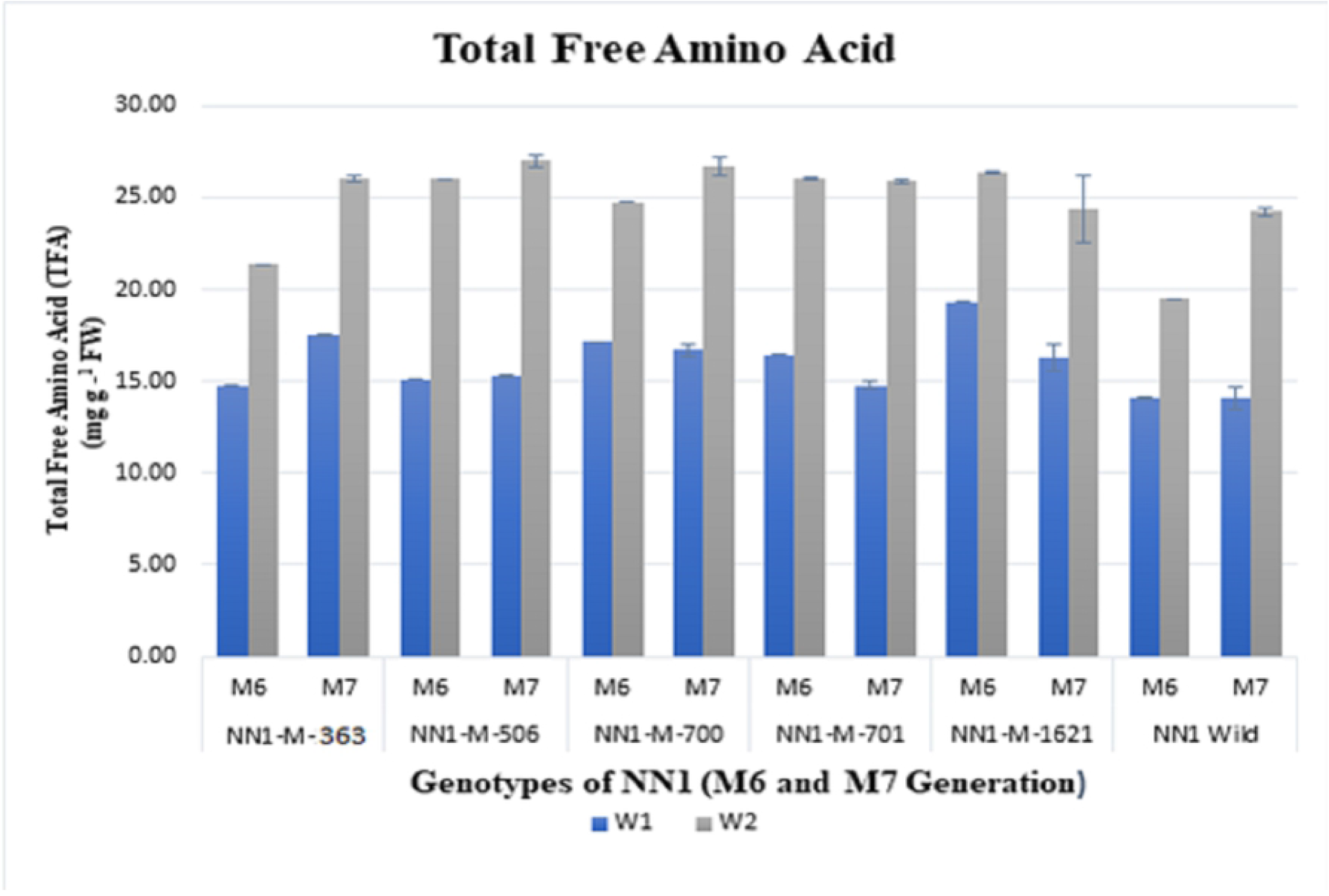
Total Free Amino Acids (TFAs) in NN-Gandum-1 genotype under well-watered (control, W_1_ regime) and rainfed (stressed, W_2_ regime) conditions. Values presented are means ± S.D of three replicates per genotype in both M_6_ and M_7_ generation. Different color indicates stress condition, blue bar for W_1_-well watered (control) and grey bar for W_2_-rainfed (stressed) conditions

#### Total Chlorophyll Contents

Drought limits the photosynthesis by retarding the synthesis of photosynthetic pigments. It is well established fact that chlorophyll contents and photosynthetic rate are positively correlated [105, 106]. In the present study, NN1-M-wild depicted 17% reduction in chlorophyll content while NN1-M-506 recorded 22% during M_7_ generation (Fig 8; Table S2). Photosynthetic pigments production reduces under drought stress because of splitting of thylakoid membranes due to desiccation of cells [107]. The drought-resistant varieties produce these pigments in abundance under drought stress [107–109]. The synthesis of chlorophyll content repressed due to water deficiency, resultantly produced ROS, and eventually caused lipid peroxidation. Physiological traits of wheat genotypes studied under rainfed conditions had high chlorophyll contents that help in improved seed production during dehydration [110–113].

**Fig 8.**
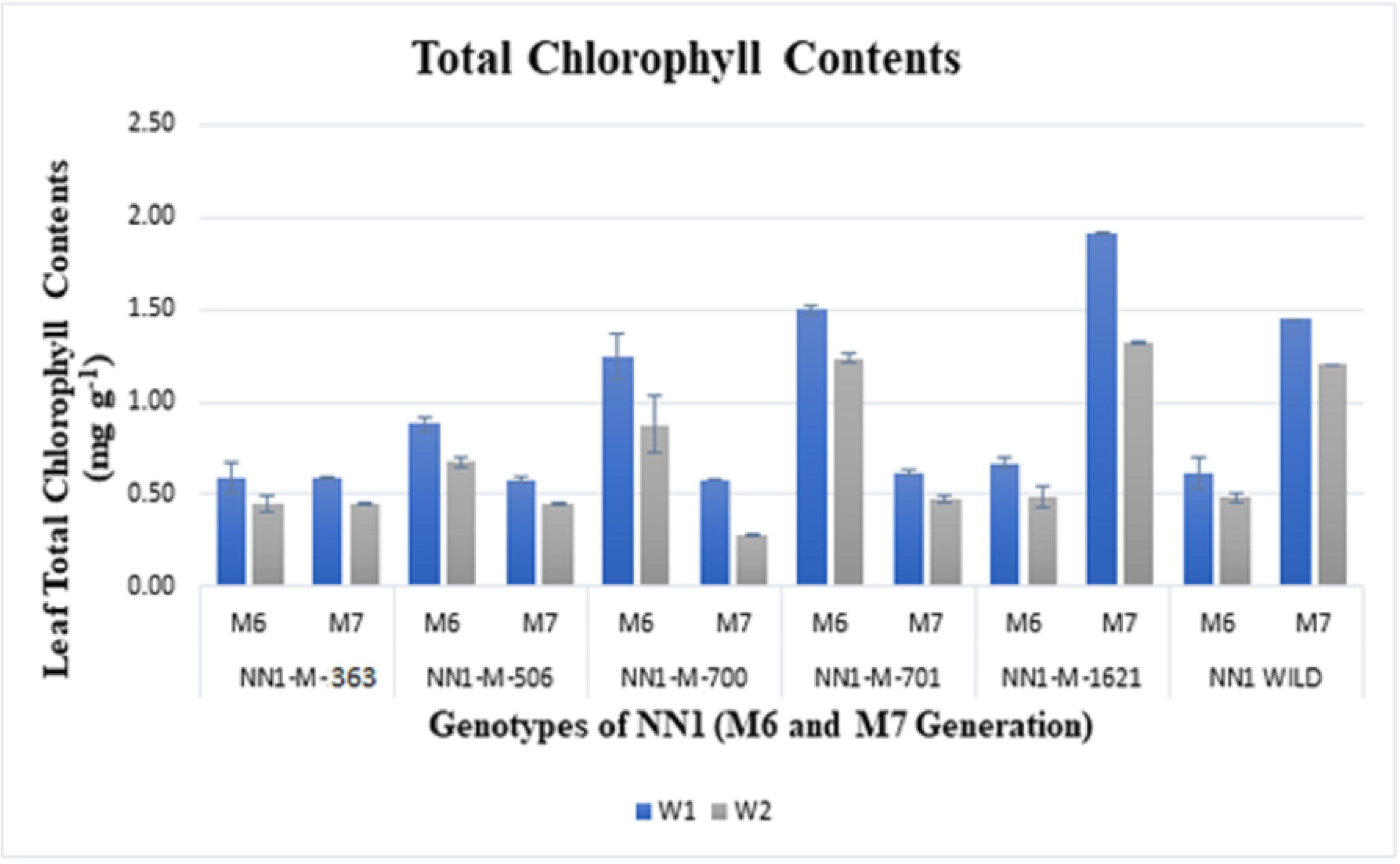
Total Chlorophyll Contents in NN-Gandum-1 genotype under well-watered (control, W_1_ regime) and rainfed (stressed, W_2_ regime) conditions. Values presented are means ± S.D of three replicates per genotype in both M_6_ and M_7_ generation. Different color indicates stress condition, blue bar for W_1_-well watered (control) and grey bar for W_2_-rainfed (stressed) conditions

#### Carotenoid Contents

A dynamic part is played by carotenoid contents to scavenge oxygen, hence, the relative quantity of carotenoid in plants approves the comparative tolerance of plant to stress. In the current experiment, maximum reduction in carotenoid content was observed in genotypes NN1-M-506 and NN1-M-701 in M_6_ generation. However, NN1-wild demonstrated 20% reduction during the first growing season while 36% reduction was recorded in the second growing season (Fig 9; S2 Table). In another study, stress tolerant varieties accumulated higher chlorophyll as well as carotenoid contents [114, 115]. The amounts of pigments reduced with constant increase in drought stress and the genotypes having increased contents of chlorophyll during dehydration are considered as drought tolerant [90, 113]. The production of carotenoid contents is affected by the genetics of the plant as well as the prevailing environments [116].

**Fig 9.**
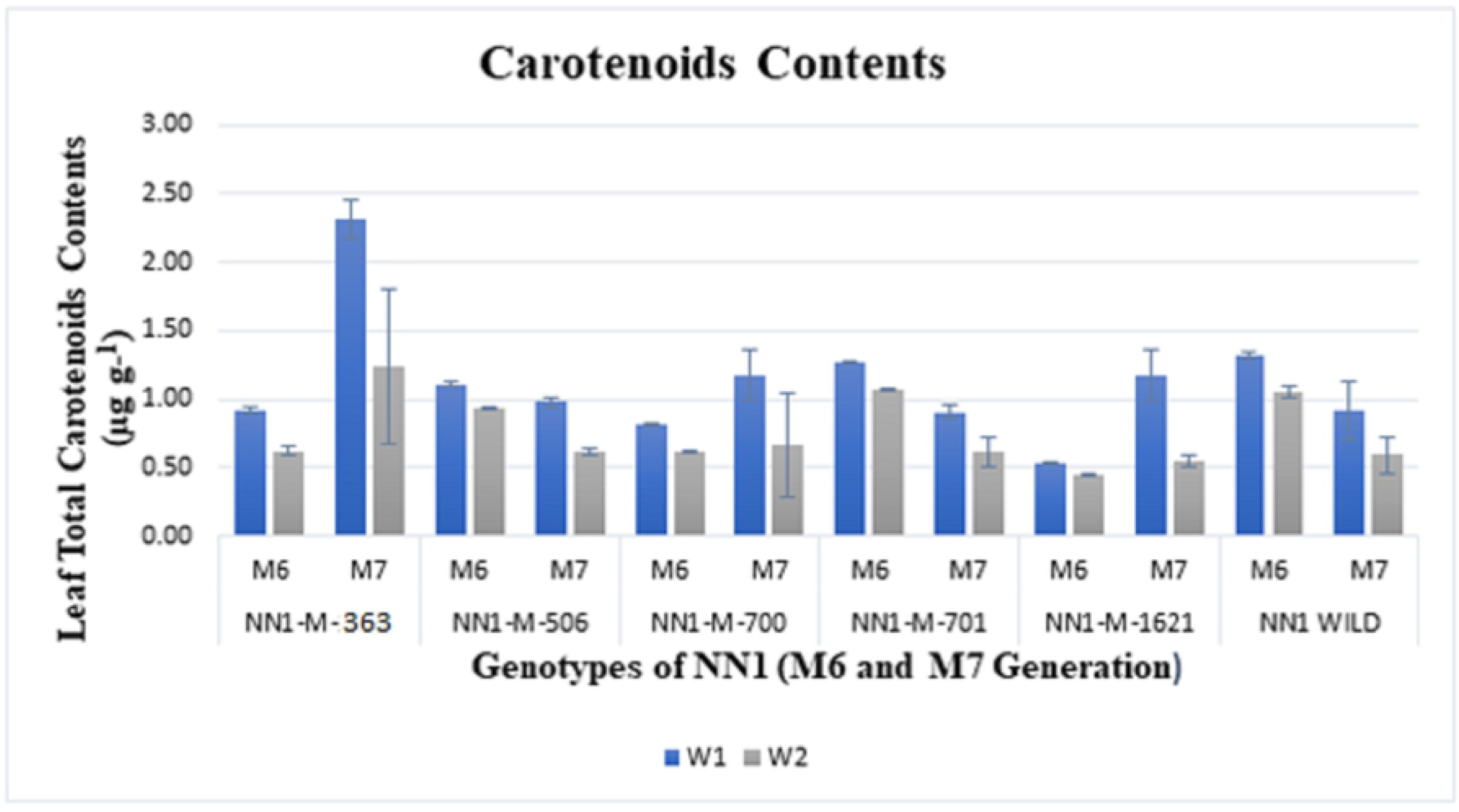
Carotenoid Contents in NN-Gandum-1 genotype under well-watered (control, W_1_ regime) and rainfed (stressed, W_2_ regime) conditions. Values presented are means ± S.D of three replicates per genotype in both M_6_ and M_7_ generation. Different color indicates stress condition, blue bar for W_1_-well watered (control) and grey bar for W_2_-rainfed (stressed) condition

#### Proline Contents

The proline synthesis escalates in response to drought stress [117–119]. Accretion of proline in wheat leaves sustains osmotic pressure of the cell [120] linked with decrease in oxidative stress and high photosynthetic ability [121–124].

In the present study, water stress induced proline contents accumulation in all mutants (Fig 10). Maximum increase (75%) was recorded for NN1-M-701, followed by 73% in each of NN1-M-506 and wild type during M_7_ generation. The wheat genotype NN1-M-701, also depicted maximum increase of 76% during M_6_ generation (S2 Table). Proline plays a vital role in membrane stability [125] as it diminishes the effects of stress by scavenging free radicals. As a signal regulator molecule, it employs many strategies that assist in mitigating drought stress [126]. Similar findings expressing the enhanced synthesis of proline under drought conditions were demonstrated in earlier studies [127, 128]. However, NN1-M-363, NN1-M-700 and NN1-M-1621 did not show significant reduction in proline contents during rainfed conditions (Fig 10).

**Fig 10.**
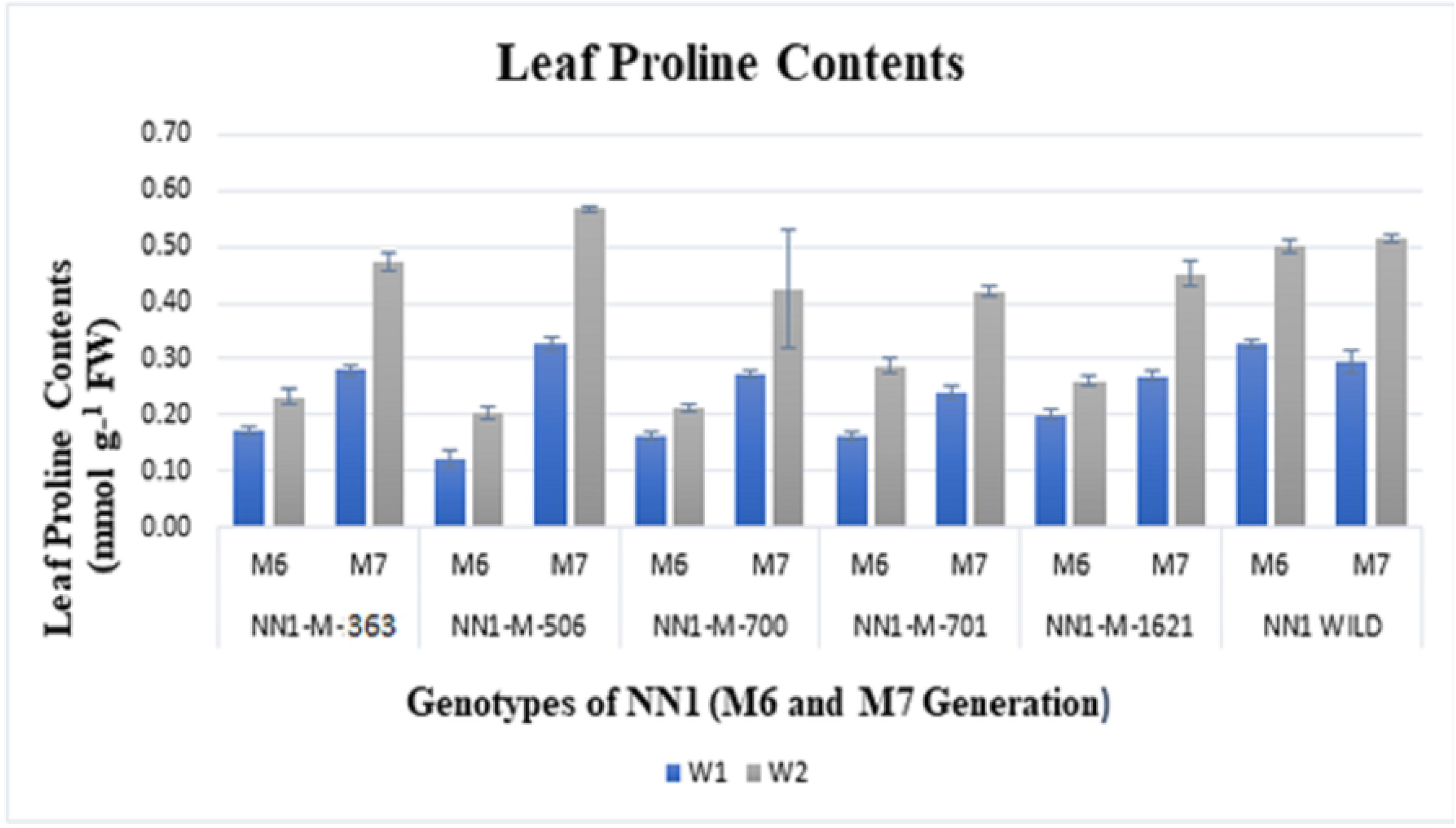
Proline Contents in NN-Gandum-1 genotype under well-watered (control, W_1_ regime) and rainfed (stressed, W_2_ regime) conditions. Values presented are means ± S.D of three replicates per genotype in both M_6_ and M_7_ generation. Different color indicates stress condition, blue bar for W_1_-well watered (control) and grey bar for W_2_-rainfed (stressed) conditions

### Superoxide dismutase (SOD) Activity

The SOD protects from the activity of ROS in plants [129]. It has the ability of affecting superoxide radicals, catalyses and converts O_2_ to O_2_, and H_2_O_2_ [28, 130, 131]. In previous experiments, an increase in SOD activity was observed in plants facing abiotic stresses, like water deficiency and toxic metal effects [132, 133]. The current results showed an increase in SOD activity in wheat mutants under rainfed conditions. In NN1-M-506 an increase of 35% in SOD activity was estimated during water stress in M_6_ generation while in M_7_ the activity was 41% (Fig 11). The SOD activity could be used as potential selection strategy for screening drought-resistant plants [134]. Thus NN1-M-506 could be used as a drought tolerant genotype in future wheat breeding experiments.

**Fig 11.**
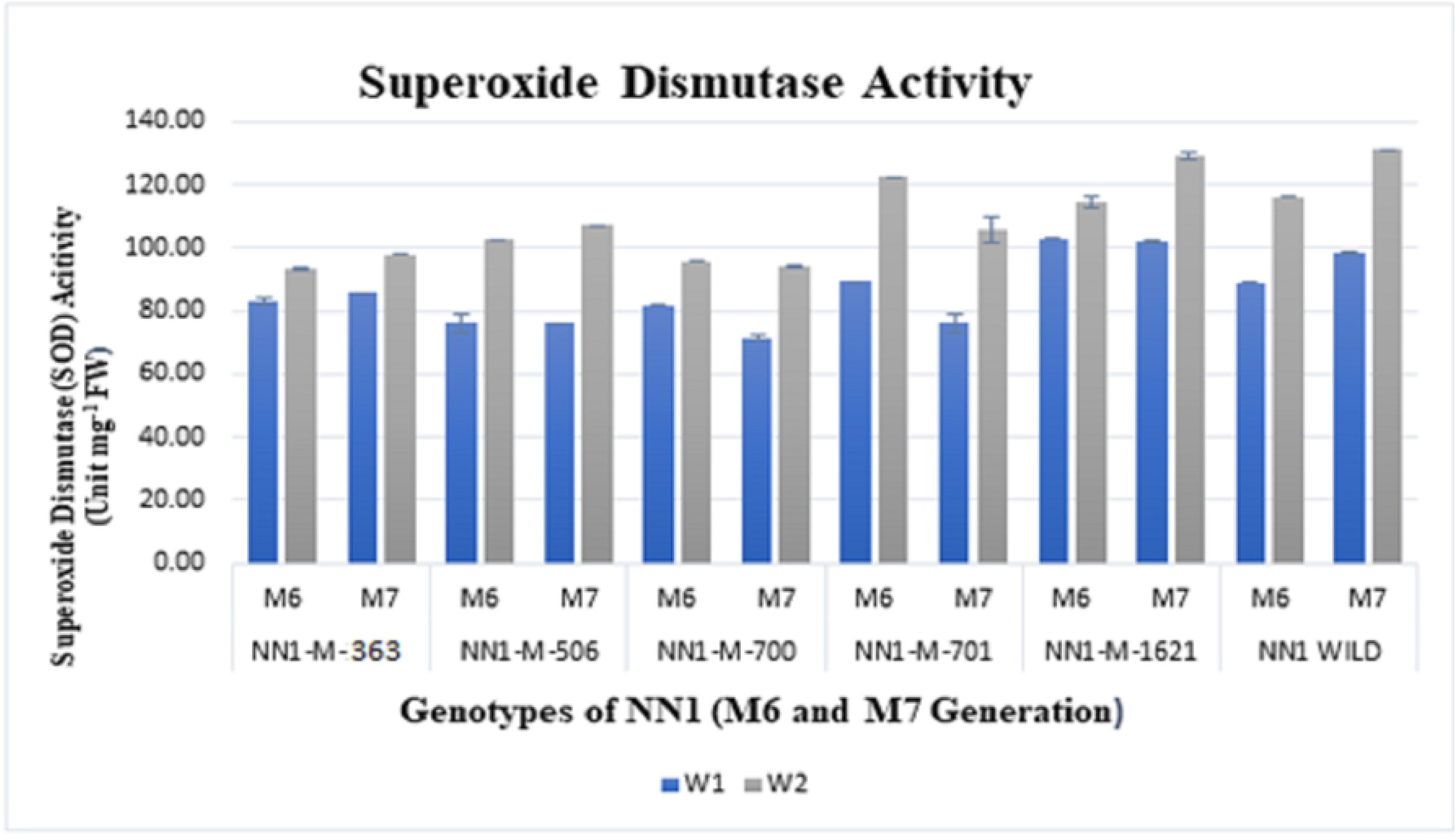
SOD activity in NN-Gandum-1 genotype under well-watered (control, W_1_ regime) and rainfed (stressed, W_2_ regime) conditions. Values presented are means ± S.D of three replicates per genotype in both M_6_ and M_7_ generation. Different color indicates stress condition, blue bar for W_1_-well watered (control) and grey bar for W_2_-rainfed (stressed) conditions

### Catalase (CAT) Activity

Catalase enzymes has an essential role in modulation of ROS in plant cells by deactivation of hydrogen peroxide (H_2_O_2_) [135]. During water limited conditions, CAT regulates harmful levels of endogenous H_2_O_2_ by catalysing a redox reaction within cell peroxisomes [28, 136]. In the current investigation, 21% and 20% increase in CAT activity was observed in NN1-M-506 and NN1-M-701, respectively under rainfed conditions in M_6_ generation (Fig 12). The same trend was observed in M_7_ for both the genotypes, confirming that these mutants could be useful source for studying drought tolerance mechanism (S2 Table). Likewise, an increase in catalase activities was noticed in wheat leaves when exposed to extreme water stress, particularly more insusceptible varieties [12].

**Fig 12.**
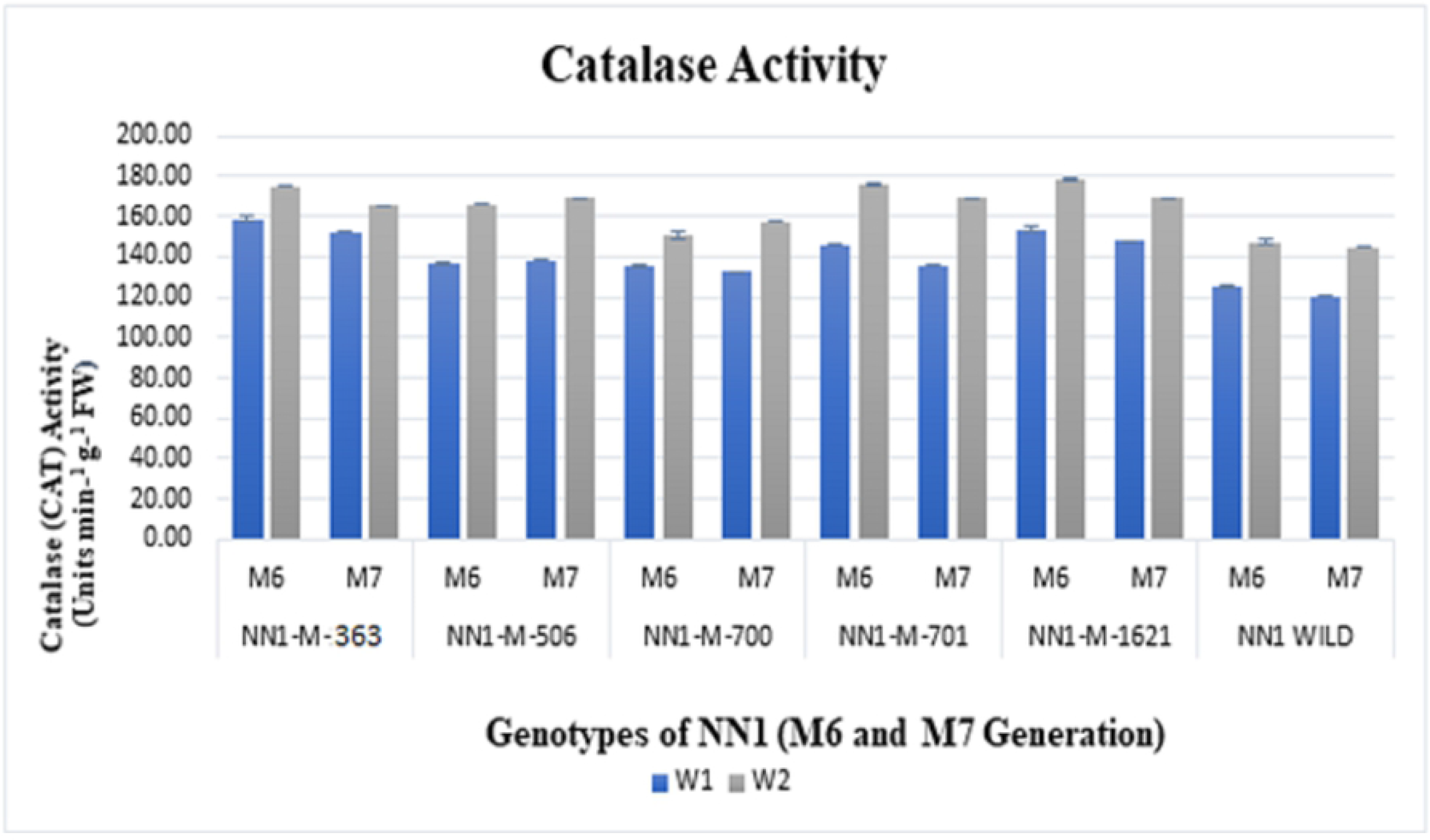
CAT activity in NN-Gandum-1 genotype under well-watered (control, W_1_ regime) and rainfed (stressed, W_2_ regime) conditions. Values presented are means ± S.D of three replicates per genotype in both M_6_ and M_7_ generation. Different color indicates stress condition, blue bar for W_1_-well watered (control) and grey bar for W_2_-rainfed (stressed) conditions

### Ascorbate peroxidase (APX) Activity

In the present study, 40% increase in APX activity was observed in NN1-M-701 in M_7_ generation while during M_6_ generation 23% increase was recorded (Fig 13). Furthermore, NN1-M-506 and NN1-M-wild also revealed improvement in APX activity. Likewise, increase in APX activity during drought stress was reported by [137]. Hence, ascorbate peroxidases are the vital enzymes in plant cells that scavenge H_2_O_2_ in different organelle of plant cells as in chloroplast and cytosol to protect against oxidative damage [130].

**Fig 13.**
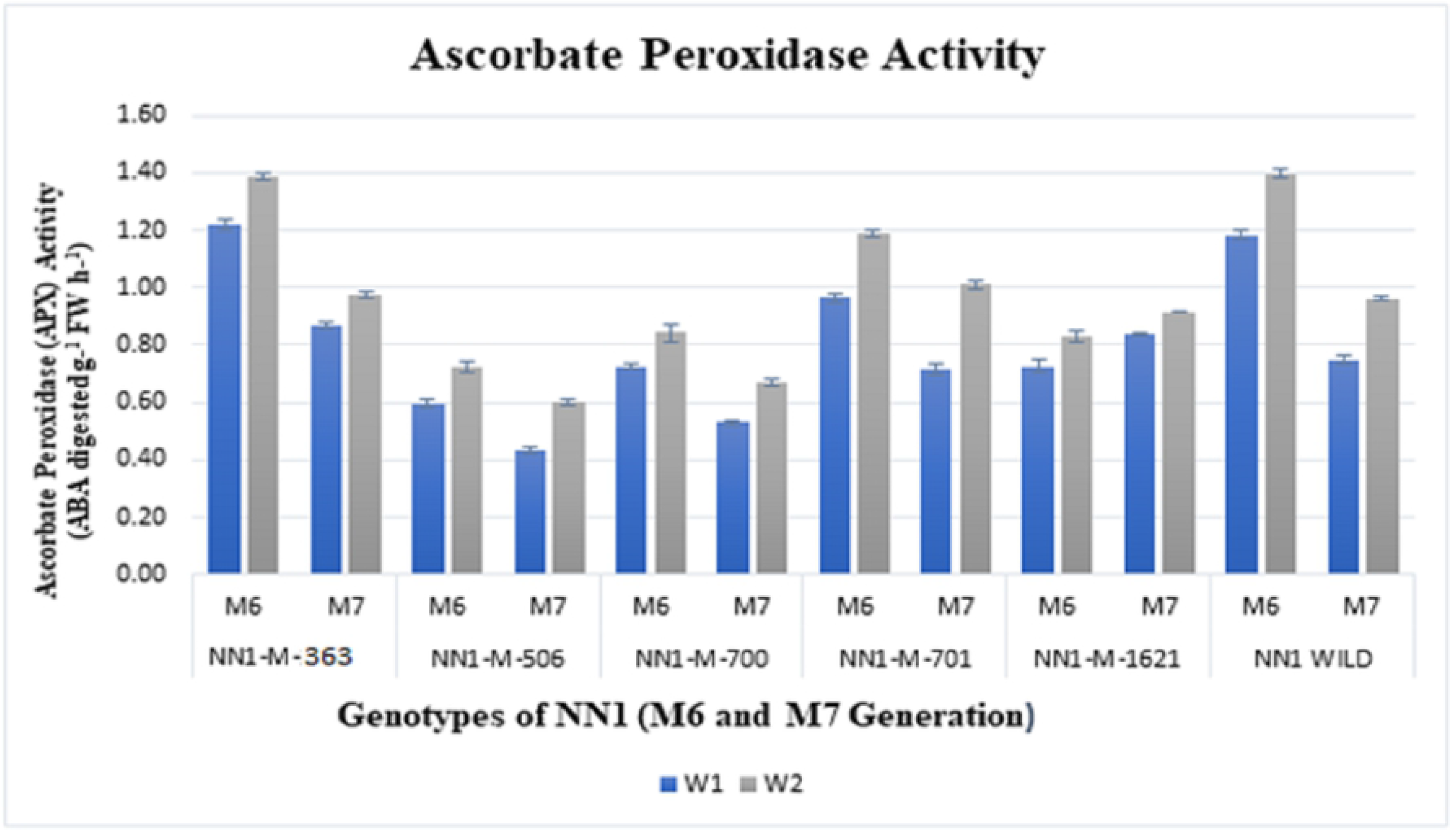
APX activity in NN-Gandum-1 genotype under well-watered (control, W_1_ regime) and rainfed (stressed, W_2_ regime) conditions. Values presented are means ± S.D of three replicates per genotype in both M_6_ and M_7_ generation. Different color indicates stress condition, blue bar for W_1-_well watered (control) and grey bar for W_2_-rainfed (stressed) conditions

### Specific Peroxiodase (POD) Activity

Reactive oxygen species production in water deficient cells results in cell damage ultimately leads towards cell death [138]. Various antioxidant systems through multiple adaptive mechanisms regulate oxidative stress, one of them is peroxidase enzyme whose activity was increased under moderate level of water deficit [31]. In the present investigations, 77% increase in peroxidase activity under limited water conditions was observed in NN1-M-701 during M_7_ generation. Owing to the rise in specific peroxidase activity during drought condition, the most affected mutant was NN1-M-700 depicting 31% increase in peroxidase activity. The NN1-M-701and NN1-M-506 were observed as highly reactive showing elevated peroxidase activity during drought stress, however, NN1-M-700 showed minimum increase in both generations (Fig 14).

**Fig 14.**
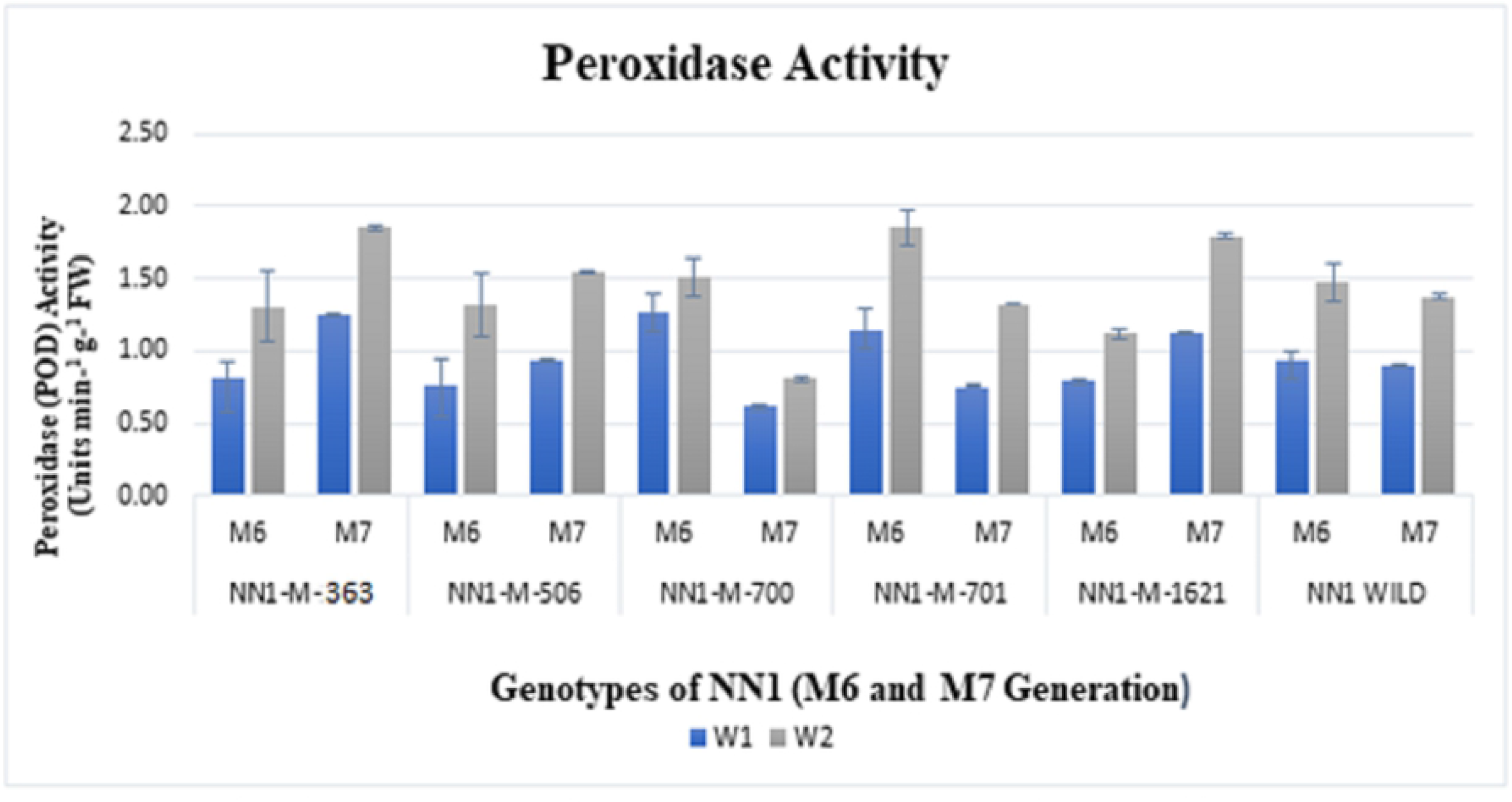
POD activity in NN-Gandum-1 genotype under well-watered (control, W_1_ regime) and rainfed (stressed, W_2_ regime) conditions. Values presented are means ± S.D of three replicates per genotype in both M_6_ and M_7_ generation. Different color indicates stress condition, blue bar for W_1_-well watered (control) and grey bar for W_2_-rainfed (stressed) conditions

The enhanced expression of peroxidase in a cell is linked with more water retention and thus rewarding tolerance to drought as it was demonstrated in *Nicotiana tabacum*. Like many other crop species, peroxidase activity was increased in wheat under water limited conditions [139, 140]. The POD is present in cytosol, vacuoles, extracellular spaces and cell walls. This enzyme is well thought stress indicator which has a wide range of selectivity for phenolic substrates and more attraction for H_2_O_2_ than that of catalase. It has the ability to utilize H_2_O_2_ in order to generate phenoxy compounds which ultimately polymerizes lignin (cell wall component) [141]. Increased POD activity revealed in the present study can be associated with the release of peroxidase localized in the cell walls [136].

### Malondialdehyde (MDA)contents

The malondialdehyde production in plants is stimulated by free radicles [142]. Membrane lipid peroxidation was evaluated with the production of MDA content that indicates the degree of membrane damage under stress. In this experiment enhancement in MDA content in all mutant genotypes was recorded (Fig 15). Maximum increase (44%) was noticed in NN1-M-701 likewise genotype NN1-M-506, also demonstrated significant increase (40%) under drought stress. A positive relationship was found among the amount of MDA and demolition of biological membranes. This suggests that increase in MDA contents, results in more lipid peroxidation and higher cell deterioration [143]. These findings have shown that under water deficit, the oxidative damage in leaves of NN1-M-701 and NN1-M-506 was greater than that of NN1-M-363, NN1-M-700 and NN1-M-1621 (S2 Table). In some other findings, higher MDA contents under water stress conditions were demonstrated, however, drought-susceptible and drought-tolerant genotypes expressed differential responses [126, 144, 145]. Moreover, the drought tolerant wheat genotype demonstrated reduction in lipid peroxidation and greater membrane stability [146].

**Fig 15.**
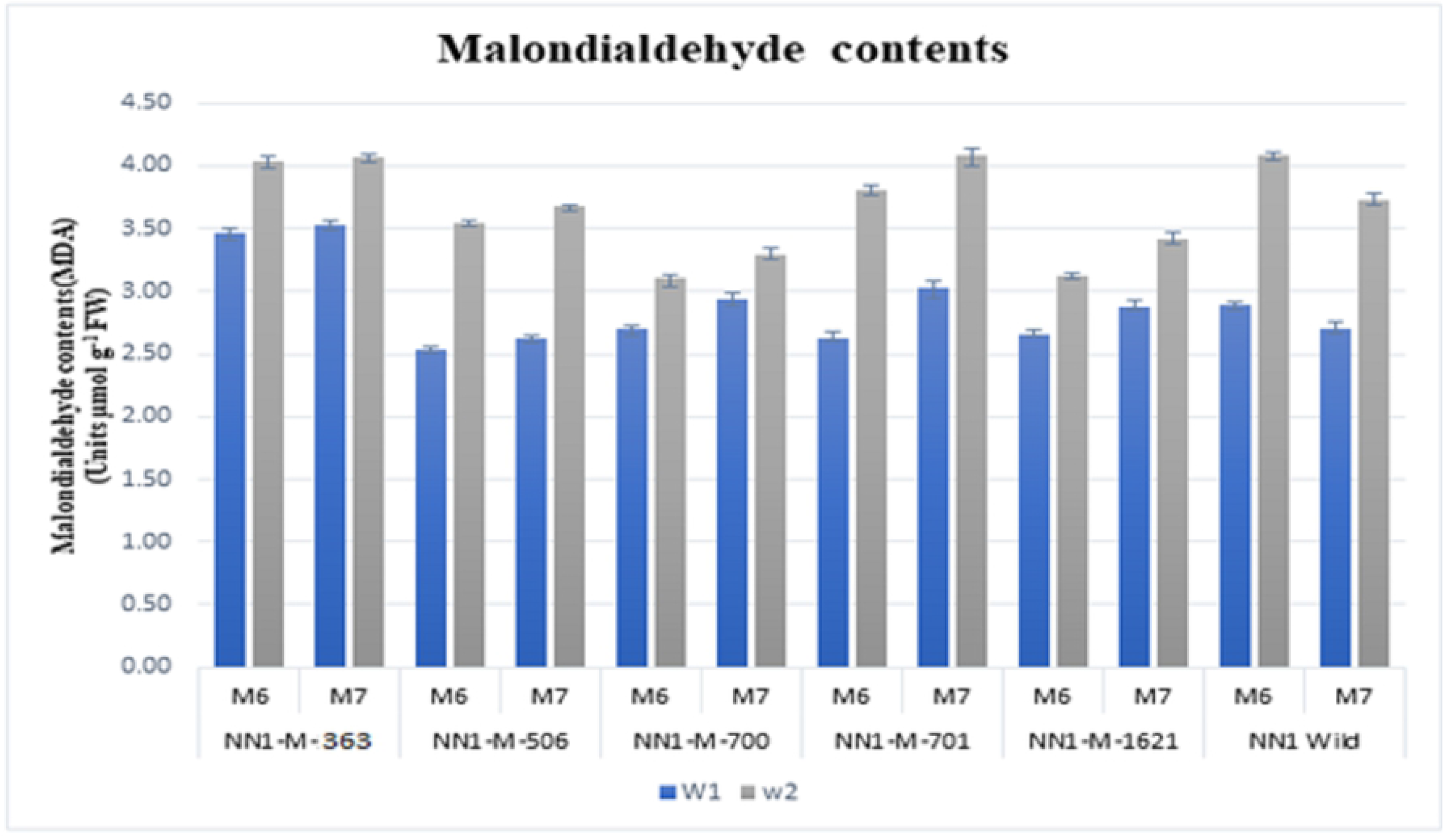
MDA contents in NN-Gandum-1 genotype under well-watered (control, W_1_ regime) and rainfed (stressed, W_2_ regime) conditions. Values presented are means ± S.D of three replicates per genotype in both M_6_ and M_7_ generation. Different color indicates stress condition, blue bar for W_1_-well watered (control) and grey bar for W_2_-rainfed (stressed) conditions

### H_2_O_2_ Activity

The H_2_O_2_ is a stress predictor of water deficit conditions in plants. It regulates respiratory pathways [147, 148]. Similarly, wheat plant protects leaves from oxidative stress by activating the antioxidant defense system. The mutant NN1-M-701 depicted the maximum increase 58% and 47% of H_2_O_2_ contents in M_6_ and M_7_ generations, respectively. Furthermore, in M_6_ generation, NN1-wild and NN1-M-506 demonstrated 49% and 53% increase in H_2_O_2_, respectively (Fig 16; S2 Table). Likewise, in M_7_ generation all the mutant genotypes displayed positive response. Multiple factors such as plant species, stress intensity and plant growth decide the H_2_O_2_ detoxification by an antioxidant enzyme [149]. During drought stress, H_2_O_2_ scavenges the ROS by activating enzymatic antioxidant defense mechanisms in wheat [150]. These findings are in close conformity that H_2_O_2_ is a stress marker that regulates the antioxidants to decrease the damage under oxidative stress [151].

**Fig 16.**
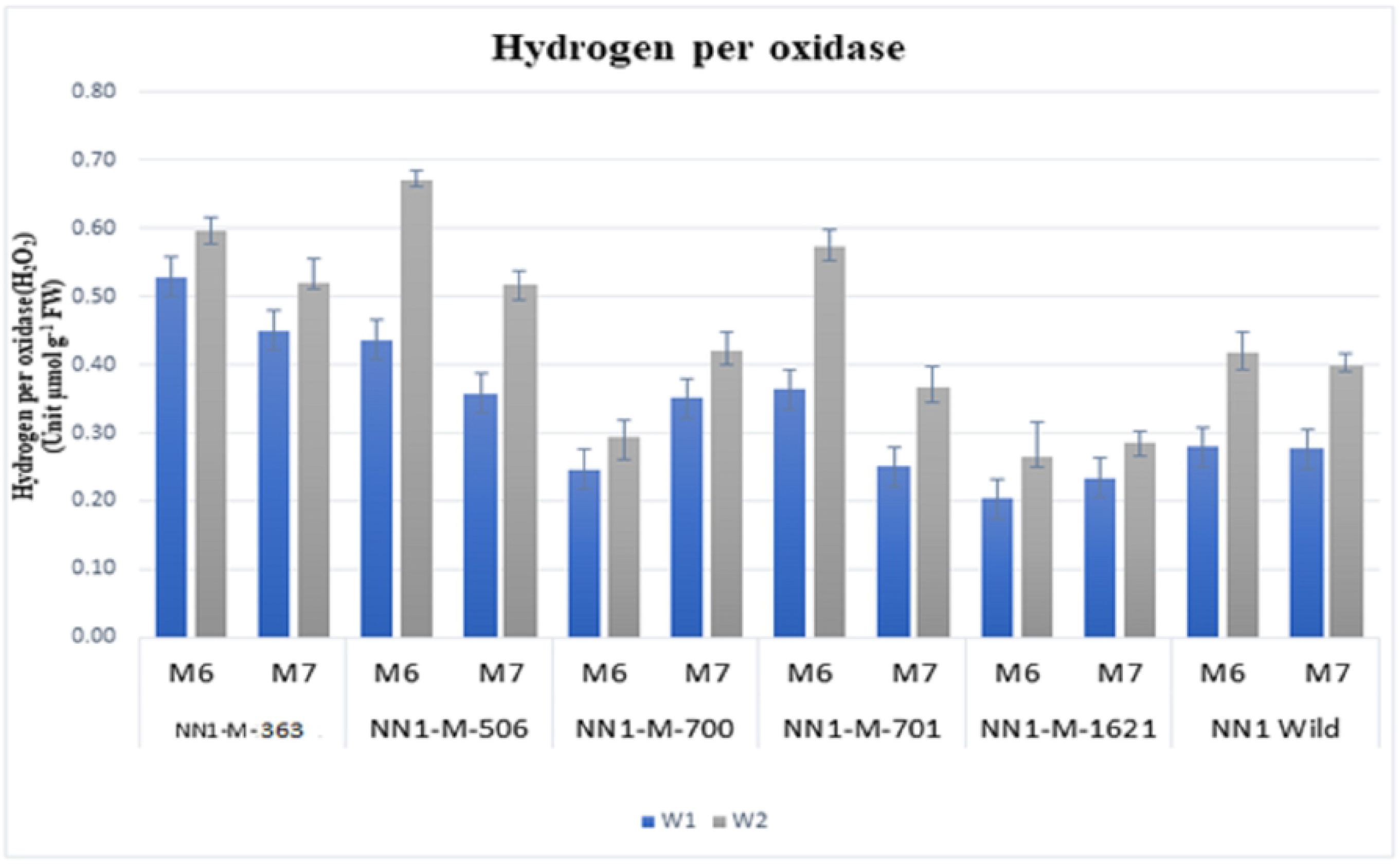
H_2_O_2_ Activity in NN-Gandum-1 genotype under well-watered (control, W_1_ regime) and rainfed (stressed, W_2_ regime) conditions. Values presented are means ± S.D of three replicates per genotype in both M_6_ and M_7_ generation. Different color indicates stress condition, blue bar for W_1_-well watered (control) and grey bar for W_2_-rainfed (stressed) conditions

## Conclusion

In the present study, it was analysed that genes relevant to agronomically significant traits can be improved via induced mutations. Also, the mutations induced in functionally important part of the gene can be identified using NGS based exome capturing assay, a strategy to save time and cost without compromising the significant mutations. This work also highlighted the distribution pattern, variation in frequency of mutation in different mutant lines of wheat. Based on the current investigations, it can be suggested that mutant wheat population NN-Gandum-1 is suitable for exploring the genetic circuits of several genetic mechanisms using forward and reverse-genetic approaches. The mutants were also explored using various biochemical and physiological assays, and showed significant variations under rainfed conditions. Hence this population can be used by the international wheat community for designing new strategies for mitigating drought stress.

## Acknowledgements

The funds for developing wheat mutant were provided by the International Atomic Energy Commission (IAEA), Vienna, Austria through a project entitled “Developing Germplasm through TILLING in Crop Plants Using Mutation and Genomic Approaches (PAK/5/047). Special appreciations are extending to Dr Cristobal Uauy, Project Leader Crop Genetics, John Innes Centre, Norwich Research Park, UK for providing lab facility for undertaking exome capture assay and analysis. I am also extremely grateful to Pakistan Agriculture Research Council (PARC) for providing funds to take this project to logical end through a project entitled “Characterization of mutants derived from EMS-derived Gandum-1 for rust and drought tolerance for sustaining wheat yield in Pakistan” (CS 049) under ALP scheme.

